# Macroscale dynamics of EEG microstates constitute the periodic and aperiodic features of the neural power spectrum

**DOI:** 10.64898/2026.07.28.741333

**Authors:** Anthony P. Zanesco, Rene A. Perez

## Abstract

The global signal characteristics of scalp-recorded electroencephalography (EEG) are composed of periodic oscillatory rhythms and aperiodic broadband fluctuations that together comprise the neural power spectrum. Spectral decomposition of these features has long served as the primary window into the macroscale characteristics of human brain activity. Here, we propose that the primary periodic rhythms and majority share of broadband spectral power within the brain’s dominant frequencies are the frequency-domain representations of the dynamics of EEG microstates. Microstates are characterized by a small repertoire of topographic voltage configurations that each reflect the momentary functional state of the cortex, and it is their time-domain dynamics that mathematically constitute periodic and aperiodic spectral features. We isolated and removed the spatial projections of distinct microstate configurations from high-density EEG using orthogonal subspace projection applied to both the surface scalp recordings and their modeled cortical generators to computationally test this dependency. Spectral parameterization of the residual power spectral density revealed that removing these spatial projections strongly attenuated alpha and theta rhythms and features of the aperiodic 1/*f* background. Selectively removing specific microstate configurations also demonstrated that each configuration aligned with spatially specific oscillatory and aperiodic 1/*f* components. Furthermore, correlational analyses of microstate temporal dynamics demonstrated that the average duration and temporal persistence of microstates strongly correlate with periodic alpha power, theta power, and the aperiodic 1/*f* background. Our findings suggest that dominant periodic and aperiodic spectral features are more accurately understood as the frequency-domain expressions of the dynamics of EEG microstates.

## Introduction

The human brain is a complex system characterized by spontaneous, moment-by-moment fluctuations in globally coordinated neural activity. Scalp-recorded electroencephalography (EEG) has long served as a non-invasive window into these macroscale dynamics. Oscillatory rhythms, such as those in the alpha band (8–14 Hz), are considered a fundamental *explanans* of brain function, generated by the coordinated activity of neuronal assemblies within cortical and thalamic networks (Buzsáki & Draguhn, 2004; Buzsáki & Vöröslakos, 2023; Fries, 2015; Varela et al., 2001). Investigating the organizing principles governing these rhythms is therefore necessary for understanding the neural basis of human cognition and behavior. Intrinsic oscillations and their resultant spectral power within narrowband frequencies have been functionally implicated in information processing, the gating of sensory inputs, and communication within distributed cortical networks (Engel et al., 2001; Foxe & Snyder, 2011; Fries, 2015; Jensen & Mazaheri, 2010). Frequency-specific EEG rhythms also correlate with established resting-state fMRI networks (Mantini et al., 2007). Yet, despite critical advances in modeling the temporal dynamics of EEG, the precise biophysical and time-domain origins of diverse oscillatory and broadband spectral features continue to be widely debated.

Historically, research has relied heavily on analyses in the frequency domain to decompose the continuous voltage time series into estimates of spectral power within functionally relevant frequencies in an attempt to isolate constituent oscillatory rhythms in the brain. Importantly, emerging perspectives challenge the assumption that the EEG signal can be adequately understood through static, time-averaged spectral analyses alone. Advances in signal processing and computational methods have facilitated attempts to better understand the non-stationary dynamics of EEG and differentiate the distinct neural phenomena that contribute to broadband aperiodic activity independent of neural oscillations (e.g., Donoghue et al., 2020; Hughes et al., 2012; van Vugt et al., 2007). This distinction is critical, as arrhythmic fluctuations and transient neural events contribute broadband power throughout the frequency spectrum. True oscillations, on the other hand, contribute power within distinct narrowband peaks rising above the aperiodic background (Donoghue et al., 2020; Ostlund et al., 2022). Temporally resolved analyses have also revealed that even genuine rhythmic oscillations are rarely sustained sinusoids and manifest instead as time- and amplitude-varying bursts of activity (Cole & Voytek, 2017, 2019; Hughes et al., 2012; Jones, 2016; Lundqvist et al., 2016; van Ede et al., 2018; van Vugt et al., 2007; Whitten et al., 2011).

The aperiodic background can be described by a 1/*f* distribution in the power spectrum forming a continuous curve with power naturally decaying at higher frequencies. The shape of this distribution mathematically reflects the fact that slower, larger-amplitude EEG fluctuations account for most of the frequency power. Modeling the 1/*f* distribution as an exponential decay function allows the aperiodic background to be parameterized by two components thought to reflect distinct neurophysiological processes (Donoghue et al., 2020; Ostlund et al., 2022): the aperiodic offset (i.e., the overall level of broadband power) and the spectral exponent (i.e., the steepness of the power spectrum’s slope). The aperiodic offset is presumed to reflect overall neuronal population spiking rates (Manning et al., 2009; Miller et al., 2014), whereas the spectral exponent serves as a putative marker of asynchronous baseline neuronal fluctuations and the excitation-inhibition balance within cortical networks (Gao et al., 2017; Preston et al., 2026). Aperiodic neural activity has been shown to have functional relevance for behavior, cognition, development, and disease (e.g., Donoghue, 2025; He et al., 2010; Hemmerich et al., 2026; Preston et al., 2025, 2026; Robertson et al., 2019).

A range of large-scale neural events are also known to contribute heavily to 1/*f* aperiodic components of the EEG power spectrum. Transient arrhythmic neural events, non-neural physiological artifacts, and time-varying bursts and non-stationary dynamics of oscillations can all contribute broadband power (Brake et al., 2024; Gyurkovics et al., 2022; Preston et al., 2026). These phenomena are sometimes considered confounds from the perspective of measuring the spectral background resulting from purely asynchronous baseline fluctuations. Alternative biophysical models, however, frame these apparent confounds as the primary constituents of the 1/*f* aperiodic components in EEG, which result from the coordinated activity of large-scale brain networks and the scaling laws underlying slower and more globally coordinated neural events (Brake et al., 2024; Buzsáki & Draguhn, 2004; Buzsáki & Vöröslakos, 2023). Understanding the origins of aperiodic spectra therefore requires investigating the extent to which large-scale neural events occurring in the time-domain constitute the aperiodic signal.

While spectral parameterizations provide insights into the macroscopic frequency composition of the EEG signal, topographic approaches are increasingly viewed as essential for characterizing the spatiotemporal dynamics of large-scale brain networks in the time domain (Michel & Koenig, 2018; Michel & Bréchet, 2026). Careful examination of the time series of voltage topographies within the multichannel EEG reveals that the scalp electrical field is momentarily organized into distinct topographic patterns that each persist for brief periods before reorganizing into new short-lasting patterns. These *microstates* result from large-amplitude co-fluctuations in the activity of distributed neural populations with durations lasting between 40 to 120 milliseconds on average (Michel & Koenig, 2018; Zanesco, 2024). A biophysical principle of EEG is that different scalp voltage topographies must be generated by the activity of different subsets of underlying cortical sources (Vaughan, 1982). The distinct topographic voltage configurations of microstates therefore reflect the activity of different collections of coordinated brain generators. As such, microstate analysis provides a time-resolved method to track the functional state of the cortex as a complement to more traditional frequency-based approaches (Michel & Koenig, 2018; Michel & Bréchet, 2026).

Empirical studies consistently show that a small repertoire of roughly four to seven highly replicable configurations explain substantial topographic variance in spontaneous EEG (Michel & Koenig, 2018; Koenig et al., 2024; Zanesco et al., 2020; Zanesco, 2024). The cross-individual ubiquity of these specific spatial patterns implies that microstates have their origins within a common electrophysiological brain network architecture in humans that is responsible for generating a considerable amount of macroscopic voltage fluctuations on the scalp. Microstates are increasingly viewed as the electrophysiological correlates of large-scale functional brain networks, with generators perhaps originating within functional networks widely identified and studied using fMRI (Michel & Koenig, 2018; Michel & Bréchet, 2026). Systematic variations in microstate configurations and dynamics are associated with individual differences in normative and clinical development (e.g., Chivu et al., 2024; Koenig et al., 2002; Takarae et al., 2022; Zanesco, 2024) and are functionally relevant for cognition (e.g., Bréchet et al., 2019; Tarailis et al., 2024; Zanesco, Denkova, et al., 2021; Zanesco, Gross, et al., 2026; Zanesco, Pandya, et al., 2026).

EEG frequency estimates originate from mathematical decomposition of time-domain voltage fluctuations. The predominance of microstates in the time-domain signal therefore makes their moment-by-moment dynamics one potential source of periodic and broadband spectral power. Microstates have long been viewed as a broadband phenomenon; however, their relationship to specific oscillatory rhythms remains contentious (Michel & Koenig, 2018; Michel & Bréchet, 2026). Indeed, microstates manifest as transient occurrences and as bursts of sustained oscillations. The time-varying dynamics of these distinct behavioral regimes introduce significant non-stationarity into the EEG signal and likely distribute power both within narrow frequency bands and across a wide broadband frequency range. Consistent with this perspective, microstates exist within delta, theta, alpha, and beta band-limited EEG, and the topographic configurations of band-limited microstates are equivalent to those identified in the broadband (Croce et al., 2020; Férat et al., 2022). Microstate durations also exhibit a heavy-tailed lognormal distribution (Creaser et al., 2021; Gschwind et al., 2015) in which prolonged occurrences ought to affect the overall shape of the broadband power spectrum by contributing power within slow frequencies (< 8 Hz) and influencing the aperiodic offset and 1/*f* exponent.

The temporal dynamics of microstates are also intimately connected to the dominant rhythms of the brain. The average durations, occurrence rates, and temporal sequences of microstates display periodicities associated with the alpha frequency band (Dziego et al., 2024; Hill et al., 2023; Javed et al., 2019; Lehmann et al., 1987; Milz et al., 2017; von Wegner et al., 2017, 2021; Wiemers et al., 2024). Information-theoretic modeling demonstrates that microstate sequence memory, durations, and transitions display rhythmic behaviors associated with periodicities in the alpha-band and its harmonics (von Wegner et al., 2017, 2021). Together, these observations motivate our hypothesis that the dominant periodic and aperiodic features of the human EEG power spectrum originate within the brain networks responsible for microstates. In support of this view, mathematical modeling has been used to argue that EEG frequency spectra and microstate dynamics are equivalent frequency- and time-domain representations of the same electrophysiological activity because the cross-spectra contain sufficient information to reconstruct microstates and their temporal dynamics (Pascual-Marqui et al., 2022).

This possibility would necessitate a reinterpretation of traditional frequency-based descriptions of EEG and expose a critical limitation in their application. Spectral estimates are typically computed channel-by-channel at the sensor level and subsequently averaged over extended spatial or temporal windows. The result is that time-averaged spectral analyses conflate the sequential activity of dozens of distinct microstates occurring each second. Spectral analyses may inappropriately aggregate activity of functionally distinct brain networks that oscillate at similar frequencies because they contribute power to overlapping scalp locations due to volume conduction. This *state-mixing* obscures the underlying spatiotemporal origins of spectra and the unique functional significance of the rapidly alternating brain networks that constitute the signal. Even time-resolved spectral methods are limited in their temporal resolution because their required integration windows can span hundreds of milliseconds (Cohen, 2014) and conflate power resulting from half a dozen or more consecutive and functionally distinct microstates. Theoretical perspectives on the etiology and functional significance of power in specific frequency bands must account for the possibility that spectral power is a time-aggregate expression of the dynamics of microstates.

In the present study, we propose that the macroscopic frequency-domain characteristics of human EEG within its dominant frequencies (< 30 Hz) result from the time-domain dynamics of microstates. We argue that the primary periodic and aperiodic components of broadband EEG spectral power are dependent on the spatiotemporal architecture of microstates and constituted by their dynamics within the scalp electric field. While prior studies have shown that microstates are broadband phenomena with alpha-band periodicities, the extent to which their time-domain dynamics constitute both periodic and aperiodic spectral features remains unknown. Addressing this requires quantifying their overall spectral contribution and determining whether the temporal dynamics of distinct microstates selectively influence these features. Accordingly, we employ orthogonal subspace projection (OSP) to isolate and computationally remove the spatial subspaces occupied by the microstate topographic configurations within the continuous EEG time series. Removing the microstate-related signals based on spatial projection of their topographic configurations maintains the continuous temporal structure of the EEG allowing for reliable frequency decomposition of the residual signal. We subsequently test their contributions to periodic and aperiodic spectra by evaluating spectral differences in the residual signal and comparing these effects against thresholds established by principal component analysis (PCA) and surrogate topographies.

OSP has direct mathematical equivalence with other established spatial filtering techniques used to unmix the EEG signal (Parra et al., 2005). OSP relies on the principle that any signal can be decomposed into orthogonal geometric components. It applies the same underlying linear algebra that is used to remove specific neural components after their identification with independent component analysis (ICA; Jung et al., 2000). Furthermore, OSP is structurally identical to signal space projection algorithms used for decades to isolate and remove the topographic variance of specific physiological artifacts (e.g., eyeblinks and cardiac artifacts; Gramfort et al., 2013; Uusitalo & Ilmoniemi, 1997). The power spectral density (PSD) of the original and residual EEG was subsequently parameterized to separate spectral estimates of periodic activity from the aperiodic background. This allowed us to computationally test the dependency of time-averaged spectral estimates on the topographic configurations of specific EEG microstates and infer how the time-domain behavior associated with these spatial configurations constitutes the overall power spectrum.

Furthermore, we empirically tested the proposed generating mechanism connecting the behavioral regimes of microstates to frequency-domain spectral estimates. Microstates often persist in time through rhythmic inversions in polarity that sustain them for several oscillatory cycles. Alternatively, they occur as polarity-fixed events with durations that tend to reflect the half-cycle frequency of the alpha band (i.e., average durations of 40–60 msec; von Wegner et al., 2017, 2021; Zanesco, 2024) but can also vary substantially with prolonged durations. We hypothesized that microstates contribute periodic alpha power when they persist in time by oscillating in polarity. Thus, greater prevalence of temporally persistent microstates should be associated with greater average alpha power in the EEG. In contrast, when polarity-fixed segments of microstates tend to be longer in duration, alpha power should be reduced because this shifts the signal’s spectral power out of the alpha band into slower frequencies. The result is a steeper aperiodic 1/*f* exponent. These temporal analyses, together with the computational removal of microstate spatial projections, aim to provide empirical support for our unified framework proposing that traditional frequency-based EEG metrics are mathematically bound to the spatiotemporal behavior of microstates and the large-scale cortical networks they represent.

## Method

### Participants

Eighty individuals (*M* age = 19.84 years, *SD* age = 4.07, age range = 18–44) were recruited from the University of Kentucky undergraduate community and participated in this study. The sample size was based on recruitment feasibility during the project grant funding period. Participants were predominately female (*n* = 62), non-Hispanic (*n* = 71), and most reported a White racial identity (*n* = 65), while the remaining participants identified as American Indian, Native American, or Alaskan Native identity (*n* = 2), Asian or Asian American (*n* = 2), Black or African American (*n* = 5), or another unspecified identity (*n* = 6). Participants verbally confirmed that they felt awake, alert, and slept at least 7 hours the prior evening (*M* = 7.83 hours of sleep, *SD* = 1.13) before the start of the session. No participants reported a history of neurological diagnoses that might interfere with interpretation or recording of EEG. Seventy-eight individuals comprised the final sample size after participant exclusions (see Procedure). The study was approved by the Institutional Review Board of the University of Kentucky. Participants provided written informed consent and received course research credit for their participation.

### Procedure

Participants sat alone at a computer monitor in a quiet, illuminated room while 128-channel EEG was recorded during the session. Participants were instructed to rest quietly, relax their face and jaw, and close their eyes for each of eight 3-minute blocks of time throughout the session. Participants were specifically instructed to remain awake, alert, and that it was “okay that their thoughts might wander to different topics and that they can think about whatever comes to mind.” An auditory tone alerted participants to the end of each block. Participants answered brief questions about their thoughts and experiences between blocks (cf. Zanesco, Pandya, et al., 2026). E-Prime 3.0 software controlled block and trigger timing information. The experimenters monitored participant compliance during the session through a half-silvered mirror in an adjacent room. Two participants were dismissed from the study and excluded from all analyses due to visible signs of drowsiness and falling asleep during the session.

### EEG Data Acquisition and Processing

EEG was continuously recorded from 128 Ag/AgCl active electrodes using a BioSemi ActiveThree system (https://www.biosemi.com/) at a sampling rate of 1024 Hz. Electrodes were distributed on the scalp according to an equiradial montage. Individual electrode locations were digitized for each participant in three-dimensions using a Polhemus Patriot digitizer (http://www.polhemus.com). EEG were processed offline using the free Cartool software toolbox version 5.06.04 (Brunet et al., 2011). Recordings were bandpass filtered between 1–40 Hz, average referenced, and downsampled to 256 Hz. EEG were visually screened for channels with intermittent connectivity or excessive periods of extreme amplitude, and these channels were interpolated using 3D spline interpolation. Individual electrode locations were also transformed into a standard 128-channel equiradial montage through the same interpolation. Each 3-minute block of resting EEG was then segmented from the original recording, concatenated into a single continuous segment, and submitted to infomax-based ICA (Jung et al., 2000) in EEGLAB (Delorme & Makeig, 2004). ICA was subsequently used to remove non-neural signal contaminants from the EEG, including sources of high-frequency noise, ocular, cardiac, and muscle artifacts. Blocks were again separated after ICA. Each block was visually inspected to identify any remaining periods of excessive noise or artifacts, which were marked for exclusion. This resulted in 620 total recording blocks of EEG (four blocks were lost during acquisition due to error). The remaining blocks were 173.19 seconds (*SD* = 5.90, range = 115–179) in length on average after excluding periods of EEG with poor signal quality or artifacts. Finally, EEG were spatially smoothed to reduce any remaining influence of outliers in the electrode montage (Michel & Brunet, 2019).

### Topographic Clustering of Microstate Configurations

A *k*-means clustering procedure determined the optimal number of representative topographic clusters of microstates by identifying the fewest number of *k* clusters accounting for the greatest global explained topographic variance (GEV) in the EEG time series (Murray et al., 2008). The clustering of microstates from EEG followed widely employed procedures that have been extensively described (see Zanesco et al., 2020; Zanesco, Gross, et al., 2026; Zanesco, Pandya, et al., 2026) but the methodological details are nevertheless summarized again in the following sections. The results of clustering are depicted in Figure 1.

**Figure 1:**
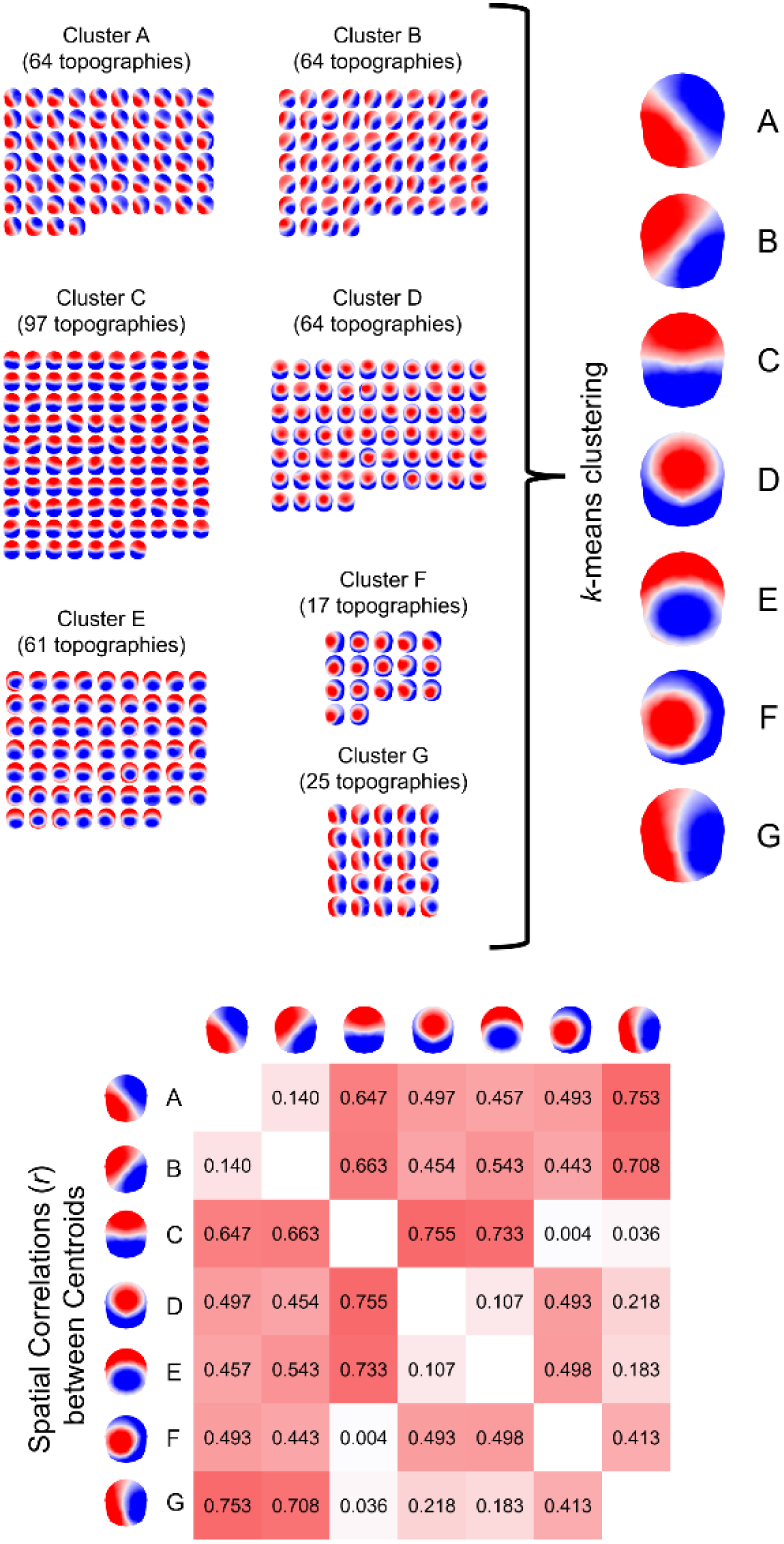
The results of topographic *k*-means clustering of 78 individuals’ EEG recordings are shown with topographic cluster centroids grouped according to the optimal *k* global clusters. Voltage maps are 2D isometric projections with nasion upwards. Seven global clusters of maps (A through G) were identified from a second round of *k*-means clustering of the individual-level centroids. These seven topographies represent the spatial voltage configurations of microstates in these data. Spatial correlations between the global cluster centroids are also provided below.

Topographic voltage maps from sets of EEG epochs at the local maxima in the global field power (GFP; Skrandies, 1990) time series were identified for clustering because these moments provide optimal representations of the topography of microstates (Zanesco, 2020). Clustering occurred separately for each individual and included all blocks of EEG from that session. Topographies were only assigned to a cluster if the spatial correlation with the centroid exceeded 0.5; and correlation values were based on the relative topographical configuration but not the polarity of the topographies by correcting the sign of the spatial correlation coefficients during cluster assignment. The optimal number of *k* clusters was selected using a metacriterion defined by seven independent optimization criteria (see Bréchet et al., 2019; Custo et al., 2017). Four to eight topographies (*M* = 5.03, *SD* = 0.82) were identified as the optimal number of *k* clusters among the 78 individuals totaling 392 topographies. The topographies explained 77.57% (*SD* = 4.59, range = 68.98–89.27) of topographic variance in individuals’ EEG recordings on average.

A second stage of *k*-means clustering subsequently identified the optimal global clusters that best explain the participant-level cluster centroids across all 78 participants. The optimization metacriterion recommended that five global clusters were the optimal *k* selection explaining 86.24% of topographic variance (i.e., GEV) among the participant-level cluster centroids. Importantly, however, we selected the seven-cluster solution to provide a more comprehensive sampling of microstate configurations previously identified in our prior work (Zanesco et al., 2020; Zanesco, Gross, et al., 2026; Zanesco, Pandya, et al., 2026) and the broader literature (Koenig et al., 2024). The seven clusters explained 88.56% of GEV and were labeled A through G according to their ordering and similarity to the meta-analytic topographic cluster solutions reported by Koenig et al. (2024). Figure 1 depicts the seven global topographic configurations alongside the 392 participant-level cluster centroids grouped according to their global cluster membership. Spatial correlations between cluster centroids (*r* range = .003–.755; see Figure 1) demonstrate the separation of centroids into distinct topographic groupings.

The microstate configurations from the four-cluster solution (spatial *r*s > .942), five-cluster solution (*r*s > .992), and six-cluster solution (*r*s > .977), are also strongly spatially correlated with specific configurations in the seven-cluster solution (see Supplementary Materials). This indicates that subspace projections using lower *k* cluster solutions would provide nearly identical configuration-specific results because the lower cluster solutions are fully encompassed by configurations in the seven-cluster solution. There was one exception in the six-cluster solution: configuration F (*r* = .776) had a somewhat lower correlation prior to the separation of configurations F and G in the seven-cluster solution. The seven configurations were also strongly correlated with centroids from the meta-analytic clusters derived from independent laboratories (Koenig et al., 2024),^1^ demonstrating the generalizability of these configurations beyond the present sample.

### Source Localization of EEG

Source estimates of the original EEG recordings and global microstate configurations were obtained using a distributed linear inverse solution (LAURA; Grave de Peralta Menendez et al., 2004) implemented in the Cartool software toolbox (Brunet et al., 2011; Michel & Brunet, 2019). The 128 electrode locations were co-registered with the Montreal Neurological Institute (MNI) template brain, and the lead field was derived from a 4-shell Local Spherical Model with Anatomical Constraints (LSMAC) head model. The average age of the sample was used to estimate skull thickness to optimize the lead field for the model. The solution space consisted of 6,926 points distributed in the grey matter of the MNI template and the inverse solution was computed without any constraints on dipole orientation (Brunet et al., 2011). LAURA accounts for depth weighting by applying local spatial autoregression that models the biophysical decay of electromagnetic fields (Grave de Peralta Menendez et al., 2004). Results were optimized with Tikhonov regularization and the L-corner criterion to automatically select the regularization parameter (Brunet et al., 2011). The 3D current density vector was obtained at each solution point. These vectors provide the estimated equivalent current dipole within each brain volume and quantify both the magnitude and orientation of the current flow through its *x*, *y*, *z* components. The dimensions of the source matrices were 20,778 (6,926 solution points × 3 vectorial orientations) × time samples.

### Computational Removal of the Spatial Projections of Microstate Configurations

We employed a spatial filtering approach utilizing orthogonal subspace projection (OSP) to selectively filter out the variance associated with specific microstate configurations in the eyes-closed resting EEG. This allowed us to isolate and remove the specific electrophysiological contributions of activity aligned with these configurations and examine their contributions to the periodic and aperiodic spectral characteristics of EEG. OSP was used to remove all microstates from the EEG, together, as well as remove each microstate separately. OSP relies on the principle that any multivariate spatial pattern can be decomposed into mutually orthogonal geometric components: a component within the linear subspace defined by a set of target spatial templates, and an orthogonal residual component. OSP was independently conducted at both the sensor level (operating on 128-channel scalp voltage potentials *X*) and the source level (operating on 20,778 vector current densities *J*). Source-level OSP acts as an anatomically weighted spatial filter that uses the biophysical constraints of the forward model to more effectively separate spatially correlated topographies than sensor-level filtering alone. Figure 2 provides an example of the results of OSP for a small epoch of EEG alongside the symbolic time series of categorized microstates for that epoch.

**Figure 2:**
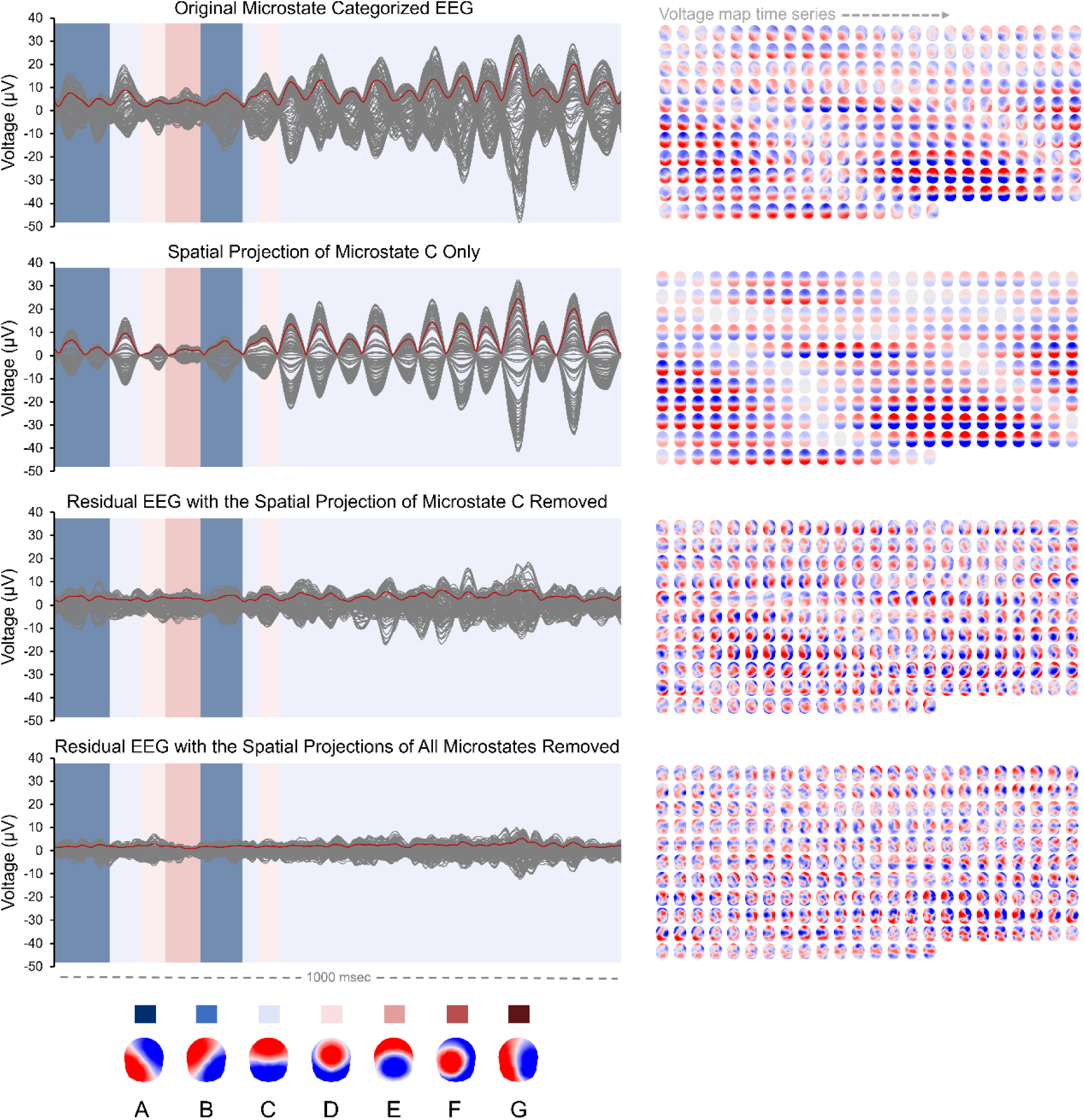
Source-level spatial filtering of the continuous EEG time series using orthogonal subspace projection (OSP) is depicted for an example epoch of a 128-channel EEG recording from one study participant. The top panel displays 1-second of the original unaltered microstate-categorized EEG alongside its 256-sample time series of voltage topographies to the right. The GFP time series is shown in red superimposed on the butterfly plot of the 128-channel EEG. The categorized microstate time series for the original data is shown in the background. The original data was source localized and forward-projected back to sensor space to serve as an equivalent baseline for the other conditions. The remaining panels show the spatial projection of microstate C isolated from the original EEG, the residual signal after the spatial projection of microstate C has been removed, and the residual signal after the spatial projections of all microstates have been removed.

#### Sensor level orthogonal subspace projection

At the sensor level, OSP was applied directly to each preprocessed EEG recording *X* (128 electrodes × time samples) using the seven group-level microstate configurations derived from *k*-means clustering. To remove all seven configurations (A–G) simultaneously, the topographies were compiled into a matrix (128 electrodes × 7 configurations) and subjected to singular value decomposition (SVD) to extract an orthonormal spatial basis *U_r_* (128 electrodes × *r*) consisting of seven (*r* = 7) 128-element spatial basis vectors with non-zero singular values. Each of the 7 vectors of *U_r_* is a topography that has been scaled to unit length and orthogonalized with respect to the other topographies. Scaling the topographies to unit length means the orthogonal subspace projection depends solely on their relative spatial configurations rather than their polarity or amplitude. The orthonormal spatial basis represents the total non-redundant information from the entire repertoire of configurations.

The orthonormal spatial basis *U_r_* was then used to geometrically project the targeted configurations out of the continuous EEG recordings. First, the continuous 128-channel EEG data (*X*) is multiplied by the transposed basis 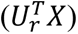. This creates a time series of activation weights that quantify how strongly the spatial patterns of the microstate configurations are present in the scalp electric field at every time sample. Second, the spatial basis is multiplied by these activation weights to reconstruct the isolated activity across the continuous EEG recording 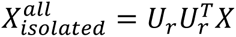. This results in a continuous EEG time series containing only voltage fluctuations aligned with the targeted configurations. Finally, to computationally remove these configurations from the EEG, 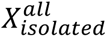 is subtracted from the original recording (*X*) to obtain the residual EEG 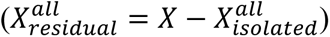. This subtraction preserves all activity that is geometrically orthogonal to the collection of microstate subspaces because they were orthogonalized with SVD. However, SVD was not required to selectively remove an individual microstate configuration. Instead, the 128-channel topographic column vector representing a single specific microstate configuration (*m*) was scaled to unit length by dividing it by its Euclidean norm to obtain the unit vector *u_m_*. The exact same three conceptual steps were then applied using this single unit vector to isolate 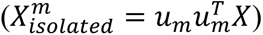 and subtract the isolated configuration projection from the original recording to generate the residual EEG 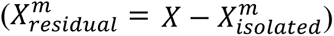

#### Source level orthogonal subspace projection

For source-level OSP, both the continuous scalp EEG recordings (*X*) and the seven group-level microstate configurations were first projected into source space using the LAURA distributed linear inverse solution. The dimensions of the resultant source matrices *J* were 20,778 (6,926 solution points × 3 vectorial components) × time samples. To remove all seven configurations simultaneously in source space, the source-localized microstate configurations were compiled into a matrix (20,778 source components × 7 configurations) and subjected to SVD to extract an orthonormal spatial basis *W_r_* consisting of seven (*r* = 7) 20,778-element spatial basis vectors with non-zero singular values. Each vector of *W_r_* represents a cortical source pattern (comprising the 3D *x*, *y*, *z* dipole components across all 6,926 solution points) that has been scaled to unit length and orthogonalized with respect to the other vectors.

The remaining steps mirrored the sensor-level projection procedure used to isolate and remove the targeted configurations out of the source-localized recordings. First, the continuous source matrix *J* is multiplied by the transposed source basis 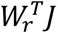 to calculate a time series of source activation weights. Second, the source spatial basis is multiplied by these activation weights to reconstruct the isolated activity across the cortical source space 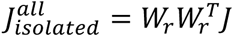. Third, 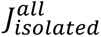 is subtracted from the original source matrix (*J*) to obtain the residual source activity 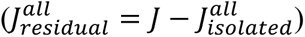. To selectively remove an individual microstate configuration (*m*) in source space, the 20,778-element source column vector for that configuration was scaled to unit length by dividing it by its Euclidean norm to obtain the unit vector *w_m_*, and the same steps were applied to isolate 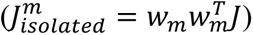 and subtract the isolated configuration projection from *J*. Importantly, because each cortical voxel comprises three directional dipole components, this projection removes only the activity aligned with the specific 3D dipole orientations of the targeted microstate subspace while preserving concurrent neural activity within the same cortical voxels that are orthogonal to that subspace.

After isolating and removing the microstate configurations in source space, the modified source matrices *J_isolated_* and *J_residual_* were forward-projected back to the 128-channel sensor space using the original lead field matrix to reconstruct the isolated and residual scalp EEG time series for spectral analysis. However, comparing the forward-projected residual signal to the original EEG is not necessarily appropriate because the inverse solution introduces spatial smoothing and might alter the amplitude characteristics of the signal. To provide an equivalent baseline for subsequent spectral analyses, the unaltered source matrices (*J*) were also forward-projected back to the scalp through the identical lead field. This ensured that the reconstructed baseline EEG, the isolated EEG, and the residual EEG all underwent the exact same spatial smoothing and attenuation from source localization prior to computing and parameterizing the power spectral density.

#### Proportion of variance explained

Finally, we calculated the proportion of variance explained by microstate configurations by examining the reduction in variance when these configurations are isolated and removed at the source and sensor-level.^2^ Specifically, we examined the variance explained directly within the 128-channel sensor space, directly within the 20,778-dimensional source space, and within the reconstructed 128-channel sensor space after forward-projecting the source-filtered signals. Variance explained was computed for the full orthogonalized subspace of all seven microstate configurations simultaneously to assess their total non-redundant contribution, as well as computed separately for each individual microstate configuration (A–G). Epochs marked as artifacts were excluded from all calculations. It is important to note that projecting out the spatial subspaces of separate microstate configurations can explain overlapping portions of the signal variance because configurations possess spatially correlated topographies. In contrast, simultaneously removing all configurations through the orthogonalized spatial basis provides the total non-redundant variance explained by the entire microstate configuration repertoire.

For projections evaluated directly at the sensor level or directly within the 3D source space, variance explained was calculated as a ratio of the isolated signal over the original signal. Algebraic subtraction of the orthogonal signal components and their forward projection through the lead field are strictly linear operations. This means summing the isolated and residual EEG (*J_isolated_* + *J_residual_* and *X_isolated_* + *X_residual_*) exactly reconstructs the baseline signal within machine precision at both the sensor and source levels. However, when the isolated and residual source matrices (*J_isolated_* and *J_residual_*) are forward projected back to the scalp, the reconstructed sensor signals can overlap because of modeled volume conduction despite being orthogonal within source space. We accounted for this in the forward-projected EEG by calculating the proportion of variance explained as the ratio of the isolated signal variance to the sum of the isolated and residual signal variances. The signal variance removed through OSP is equivalent to the total reduction in the area under the curve of the power spectral density (Cohen, 2014).

### Spectral Power Estimation and Parameterization

The power spectral density (PSD) was calculated within artifact-free epochs of the eyes-closed resting EEG using Welch’s spectral density estimation method using a 4-second Hanning window with 50% overlap (Welch, 1967). The PSD was obtained for the original 128-channel recordings as well as the forward-projected isolated, residual, and reconstructed original EEG. The estimated PSD was subsequently parameterized using the *specparam* toolbox (Donoghue et al., 2020; Ostlund et al., 2022) to separate genuine neural oscillations from the broadband aperiodic background and estimate the aperiodic 1/*f* offset and exponent. The *specparam* algorithms model the PSD as a combination of aperiodic components and distinct periodic peaks. Spectral parameterization was applied to the average PSD of each electrode across a fitting range of 2 to 38 Hz. Periodic components were identified using a minimum peak height threshold of 0.1, a maximum of 4 peaks, and peak width limits of 1–8 Hz. The candidate peak detection threshold remained at the default of 2 standard deviations. The aperiodic component was modeled using the fixed mode without a knee parameter. For each electrode, the peak power (height above the aperiodic fit), center frequency, and bandwidth were obtained for the most prominent peak identified within the theta (4–8 Hz) and alpha (8–14 Hz) frequency bands. The aperiodic offset (the overall broadband level or intercept) and the aperiodic exponent (the steepness of the 1/*f* slope) were also obtained for each electrode. Finally, these electrode-level parameters were averaged across the entire montage to obtain a global, electrode-averaged value for each periodic and aperiodic spectral measure. Pooled *R*^2^ was calculated for each recording to evaluate the overall fit of the *specparam* models across all electrodes by summing the residual and total sums of squares from models for each electrode and subtracting the ratio of the total residual variance to the total variance from one.

### EEG Dimensionality and Surrogate Comparisons

We established several comparison thresholds for variance explained by the seven microstate configurations. Specifically, we obtained the mathematically optimal spatial basis of the EEG signal using principal components analysis (PCA) and compared our results to OSP of surrogate-generated smoothed noise topographies.

#### Dimensionality and eigenvalue decomposition

PCA was conducted on each of the original 128-channel recordings (*X*) and their corresponding 20,778 source current density components (*J*) to evaluate the spatial dimensionality of the eyes-closed resting EEG. The eigenvalues from PCA were averaged across all individuals to obtain the grand average proportional and cumulative variance explained by each rank-ordered spatial component. The effective dimensionality was defined as the minimum number of components required to explain 95% of the grand average cumulative variance. This threshold establishes the theoretical minimum number of spatial topographies required to account for 95% of the signal variance. The cumulative variance of the first seven principal components was also evaluated to establish the absolute maximum variance that can be explained by any seven possible topographies. The eigenvectors representing the topographies of those seven PCA components explain more variance in the EEG than any seven possible spatial topographies. This includes the global microstate topographies identified through clustering. Finally, the eigenvectors representing the spatial topographies of the first seven components were extracted from the 128-channel recordings and visualized using the mean spatial covariance matrix across all individuals (see Supplementary Materials).

#### Spatially smoothed surrogate topographies

A surrogate analysis was conducted using topographies generated from spatially smoothed noise. A total of 10,000 independent topographies were randomly generated. Each topography consisted of 128 values sampled from a standard Gaussian distribution (µ = 0, σ = 1). A spatial smoothing filter was applied to the noise to generate patterns resembling the broad scalp topographies of actual EEG that result from volume conduction. A *k*-nearest neighbors algorithm identified the five closest neighboring electrodes for each electrode based on the electrode coordinates. Each value at an electrode was replaced with the mean of its own value and its five nearest neighbors. This local averaging process was applied three times for each topography, resulting in 10,000 spatially smoothed surrogate topographies (see Supplementary Materials for depictions of the surrogates).

Collections of these topographies were removed from the 128-channel eyes-closed resting EEG using OSP. Seven topographies were randomly sampled without replacement from the 10,000 surrogates for each of 5,000 permutations. This maintained the same dimensionality as the microstate configuration subspace for each permutation. Each set of seven random noise topographies were orthogonalized with SVD and subsequently removed from the EEG using OSP. The percentage of variance explained by the spatially smoothed noise topographies was obtained for each of the 5,000 permutations to construct a null distribution of explained variance for each EEG recording. The residual EEG was reconstructed after removal of topographies using the specific random permutation that explained variance closest to the mean of each recording block’s null distribution. Finally, the PSD of the residual EEG was calculated and parameterized using the same methods described above to compare to the microstate configurations.

The surrogate comparison was further extended to the source domain. The 10,000 spatially smoothed sensor-level noise topographies were source localized using the same procedure described above. We identified the specific random permutation of seven topographies that captured a percentage of variance closest to the mean of the sensor-level null distribution for that recording block. These seven source topographies were orthogonalized with SVD and removed from the source-level EEG using OSP. Finally, the forward-projected residual EEG was reconstructed, and the PSD was calculated and parameterized.

### Microstate Categorization and Lognormal Distribution of Microstate Durations

To evaluate the temporal dynamics of microstates and their associations with periodic and aperiodic spectral features, we categorized microstates within the continuous EEG recordings according to standard procedures (see for example Zanesco et al., 2020; Zanesco, Gross, et al., 2026; Zanesco, Pandya, et al., 2026) and quantified the durations of microstate occurrences. Each time series sample of the multichannel EEG was categorized according to the seven global configurations based on its spatial correlation. Polarity was ignored during categorization by correcting the sign of the spatial correlation coefficient. This resulted in a symbolic time series of categorical labels describing sequences of microstates. EEG samples that correlated poorly with global configurations (spatial *r*s < 0.5) were left uncategorized. Additionally, temporal smoothing was applied to the symbolic sequence (Besag factor *k* = 10 and *b* = 2; Pascual-Marqui et al., 1995) and by ignoring assigned microstates that were present for 6 or fewer consecutive samples (i.e., ≤ 23 msec) and instead splitting the time points between the preceding and subsequent microstates in the series. On average, 96.27% (*SD* = 3.01, range = 67.82–99.65) of time series samples from the 620 total recording blocks were categorized according to the seven global microstate topographic configurations. The seven microstate configurations in total explained 61.75% (*SD* = 5.65, range = 48.10–75.69) of the global explained topographic variance in the EEG time series on average.

Microstate durations were calculated from the categorized microstate time series using two approaches inclusive of microstates oscillating in polarity, as well as separately calculating durations for fixed-polarity occurrences that manifest as isolated transient events or as half-cycle segments within continuous polarity inversions of the same microstate. In the first polarity-independent calculation, successive topographies assigned to the same microstate configuration were evaluated as single, continuous occurrences regardless of the polarity of the time sample’s topography. In the second polarity-fixed calculation, we evaluated microstate durations while accounting for the polarity of the topography by considering the sign of its spatial correlation with the global microstate configuration that categorized the time series samples. This resulted in continuous microstates being segmented into their constituent phases by isolating the durations of the half-cycle segments as well as the transient microstates that occur without successive inversions in polarity of the same topographic configuration. Unlabeled sections of data were ignored and fixed-polarity segments lasting fewer than 5 samples (< 19.5 msec) were excluded so that only uninterrupted segments were quantified.

Figure 3 provides a depiction of the time-varying dynamics of microstates and their transient and oscillatory behavior. Several microstates are depicted occurring prior to the extended occurrence of microstate C oscillating in polarity. In the first (polarity-independent) approach for calculating microstate durations, the extended occurrence of microstate C is evaluated as a single, continuous event successively inverting in polarity for more than 250 msec. In the second (polarity-fixed) approach, the sustained oscillation is subdivided into five distinct half-cycle segments (ranging from 43 to 59 msec each) separated by polarity inversions. The duration of each segment is shown alongside the topographic voltage configurations at the GFP local maxima.

**Figure 3:**
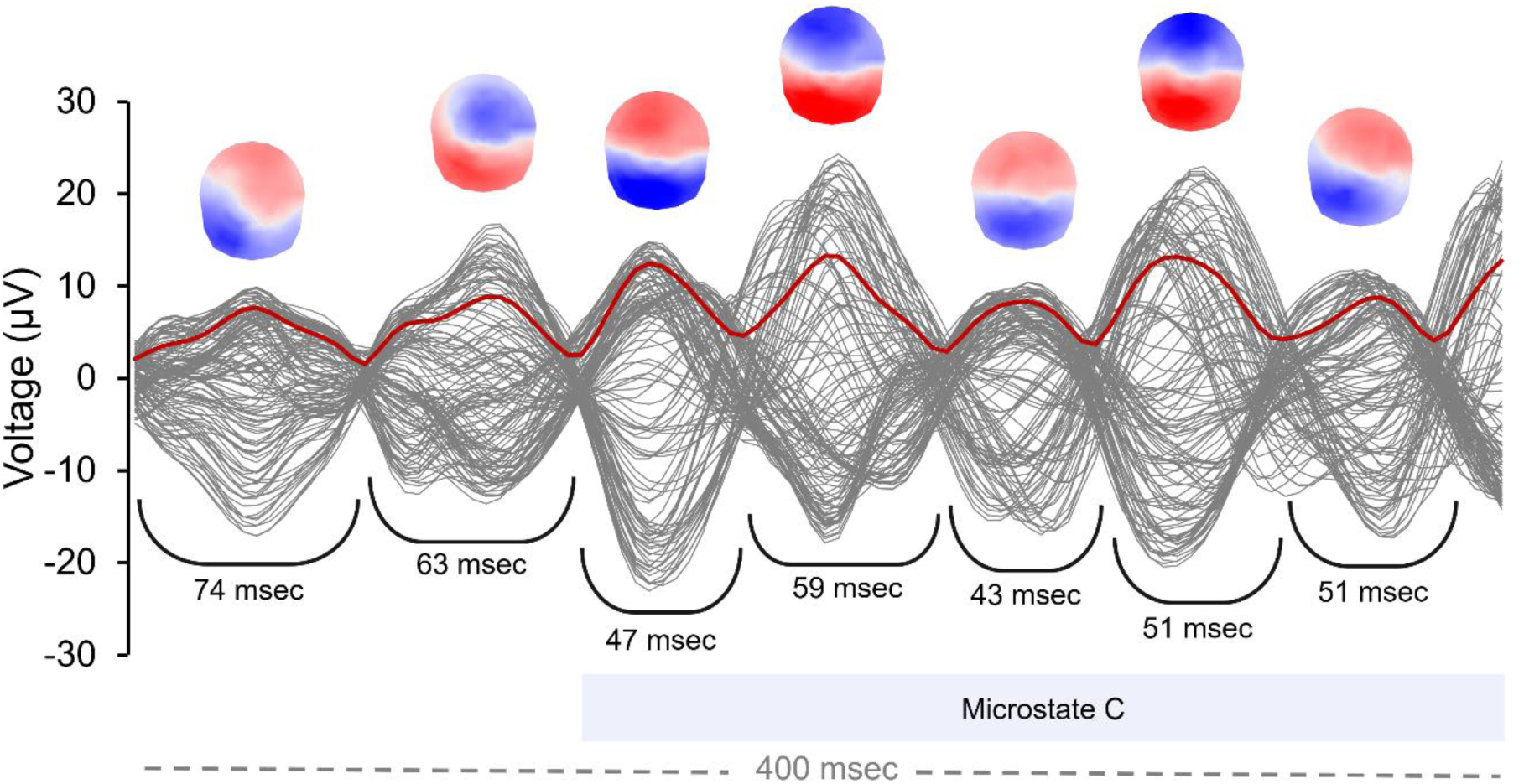
The butterfly plot depicts the transient and oscillatory behaviors of EEG microstates. The GFP time series is shown in red superimposed on the voltage waveforms of 128-channel multichannel EEG (gray lines). Topographic voltage configurations are shown at the local GFP maxima and depict microstate C oscillating in polarity over an extended duration (> 250 msec). The polarity-fixed half-cycles of the entire oscillatory event range from 43 to 59 msecs in duration as indicated by each segment in brackets, identifying this behavior as an oscillation in the alpha (8–14 Hz) frequency band. The two initial transient microstates (categorized here as microstates A and D, respectively) have longer durations than the half-cycle segments of microstate C but do not oscillate continuously in polarity. Polarity-independent calculations of microstate duration treat the entire microstate C envelope as a single continuous event (shown as the light blue bar). Polarity-fixed calculations instead evaluate the durations of the individual half-cycles once the oscillation is divided into its constituent half-cycle segments based on polarity inversions.

Microstate durations are strictly bounded and exhibit a characteristic heavy-tailed rightward skew in their distribution (Gschwind et al., 2015). As such, the arithmetic means and standard deviations of microstate durations are biased representations of the central tendency and variance of the distribution. Therefore, we obtained the lognormal location parameter (μ) and the lognormal shape parameter (σ) describing the heavy-tailed distribution of microstate durations by applying a natural log transformation to the microstate duration values (in seconds) and directly computing the mean and standard deviation of the log-transformed values. These values were calculated for each recording block within individuals across microstate configurations (A–G), as well as considering all microstate configurations together. The location parameter (μ) reflects the central tendency of the durations in log values, while the shape parameter (σ) quantifies the variance of the lognormal distribution. Importantly, the shape parameter explicitly quantifies the extent of the heavy right tail of the distribution because microstate durations are bounded and right skewed. A larger lognormal µ represents greater median durations of microstates, whereas a larger lognormal σ represents more prevalent heavy-tailed durations of microstates that stretch far beyond the median duration. These parameters were separately extracted for both the polarity-independent and polarity-fixed distributions to serve as variables in subsequent analyses.

### Analysis

Multi-level mixed-effects models were used to evaluate the impact of separately removing microstate topographies at both the sensor level and the source level on the macroscopic periodic and aperiodic features of the EEG. Mixed-effects models were also used to estimate the mean proportion of variance explained and 95% confidence intervals for the microstate configurations, spatial principal components, and surrogate topographies. Models were estimated with restricted maximum likelihood using the PROC MIXED procedure in SAS 9.4. The multiple observations from recording blocks were nested within individuals, and random intercepts for participants were included in the models. Degrees of freedom were estimated using the Kenward-Roger approximation. To account for heteroscedasticity across the different microstate configurations, residual variances were allowed to vary by microstate configuration.

The periodic peak power, center frequency, and bandwidth, as well as the aperiodic offset and aperiodic exponent, were compared between residual and original EEG for the average across all electrodes. Electrode-averages for each recording block were calculated across all electrodes that had an identifiable periodic or aperiodic component within a specific projection condition and were only included in analyses when a periodic peak or aperiodic component was identified in a minimum of 10 electrodes. This ensured that the electrode-averaged parameters reflected globally robust spectral activity. The consequence, however, was varying numbers of valid recording blocks across individuals. Missing data was accounted for in our analyses because restricted maximum likelihood estimation accommodates missing and unbalanced data. Although center frequency and bandwidth were calculated and examined (see Supplementary Materials), our primary analyses focused on periodic peak power, aperiodic offset, and aperiodic exponent. The spatial filtering condition was entered as a categorical fixed effect in models to compare the original EEG (i.e., the intercept) to the eight residual signals: the signal with all microstates removed, and the signals with configurations A through G removed individually. For the primary hypothesis tests evaluating electrode-average comparisons, a conservative significance threshold of *p* < .001 was applied to control the overall family-wise error rate. To explore the spatial distribution of these effects for periodic peak power, aperiodic offset, and aperiodic exponent, we compared the residual and original EEG for each separate electrode. We adjusted the significance threshold of these comparisons by adapting Nyholt’s (2004) method to control the family-wise error rate across all 128 electrodes. This approach determines the effective number of independent comparisons by accounting for the large statistical dependency among electrodes. The threshold for significance was α = 0.0016 for aperiodic-adjusted alpha, α = 0.0016 for aperiodic-adjusted theta, α = 0.0006 for the aperiodic offset, and α = 0.0011 for the aperiodic exponent. For electrode-level analyses, the absence of an identified periodic or aperiodic component at a specific electrode during a given recording block was treated as a missing observation.

Additional mixed-effects models evaluated associations between the temporal dynamics of microstates and the periodic and aperiodic spectral features of the EEG. The lognormal parameters (location µ and shape σ) of microstate durations served as dependent variables. Separate models evaluated the lognormal parameters from polarity-independent and polarity-fixed duration distributions. Observations were nested within individuals and random intercepts included for participants. Residual variances varied by microstate configuration. The global, electrode-averaged spectral features (e.g., alpha and theta peak power, aperiodic offset, and aperiodic exponent) were entered as continuous fixed-effect predictors, alongside the categorical fixed effects of microstate configuration (all configurations and, separately, configurations A–G). Simple slopes were estimated from the interaction between spectral features and configuration to estimate the specific association between each spectral feature and duration parameter for all microstates together and each separate configuration. To contextualize the magnitude of these associations, parameters were averaged across recording blocks for each participant and evaluated using correlation coefficients (*r*) and ordinary least squares quadratic regression.

## Results

To evaluate the contribution of microstate dynamics to periodic and aperiodic spectral features, we parameterized these features of the PSD and compared the original EEG to the residual EEG following the targeted removal of the spatial subspaces of microstate configurations at the source- and sensor-level through OSP.

### Removing the Spatial Projections of Microstates Attenuates Macroscopic EEG Power

We first evaluated the proportional reduction in total variance, equivalent to the total reduction in area under the curve of the PSD, following source- and sensor-level OSP. This quantified the effect of removing the microstate configurations on the global signal (see Table 1). Simultaneously removing the spatial projections of all seven microstate configurations resulted in profound attenuation of the source and sensor-level EEG signal. Removing the microstate configurations directly at the sensor level explained 90.91% (95% CI [90.39, 91.43]) of the total variance. Prior to forward projection, the seven microstate configurations explained 44.94% (95% CI [44.24, 45.65]) of the variance among the distributed vector current densities at the source level. After source-level removal of these configurations and forward-projection of the EEG back to the scalp, the microstate configurations accounted for 82.71% (95% CI [82.01, 83.42]) of the total variance in the reconstructed 128-channel EEG. Additionally, there were variable contributions to variance explained across configurations (see Table 1). Microstate configuration C accounted for the largest share of variance at both the source and sensor-levels. Microstates A, B, D, and E explained moderate amounts of variance, whereas configurations F and G explained the least amounts of variance.

**Table 1:** Model-estimated mean variance explained and 95% CI through OSP removal of microstate configurations.

|  | Sensor OSP in Sensor EEG | Source OSP in Source EEG | Source OSP in Sensor EEG |
|---|---|---|---|
| All removed | 90.9118 [90.3922, 91.4314] | 44.9442 [44.2433, 45.6452] | 82.7149 [82.0139, 83.4159] |
| Microstate A removed | 30.1925 [29.6007, 30.7844] | 11.9496 [11.3721, 12.5271] | 33.1016 [32.3283, 33.8749] |
| Microstate B removed | 33.2604 [32.6567, 33.8642] | 13.8612 [13.2568, 14.4657] | 36.0629 [35.2670, 36.8589] |
| Microstate C removed | 53.4594 [52.7724, 54.1465] | 21.5004 [20.8642, 22.1366] | 52.4169 [51.5809, 53.2529] |
| Microstate D removed | 36.0817 [35.4184, 36.7451] | 17.0046 [16.3904, 17.6187] | 30.6411 [29.8257, 31.4564] |
| Microstate E removed | 34.6609 [34.0157, 35.3062] | 11.4202 [10.8158, 12.0246] | 28.1170 [27.2876, 28.9464] |
| Microstate F removed | 13.8135 [13.2149, 14.4121] | 9.5385 [8.9521, 10.1249] | 16.0083 [15.2496, 16.7669] |
| Microstate G removed | 15.2472 [14.6324, 15.8620] | 6.1616 [5.5822, 6.7410] | 16.6815 [15.9156, 17.4474] |

Note: Mean proportion of variance explained (%) is provided after the removal of microstate configurations within the 128-channel sensor space (Sensor OSP in Sensor EEG), directly within the 20,778-dimensional source space (Source OSP in Source EEG), and within the reconstructed 128-channel sensor space after forward-projecting the source-filtered signals (Source OSP in Sensor EEG). Mean estimates are provided from mixed-effects models based on the fixed effects of microstate from 78 participants and 620 blocks of EEG. 95% confidence intervals (CI) accompany the model estimated means.

#### Principal component analysis (PCA) and effective dimensionality

PCA confirmed the expected low dimensional structure of the EEG and established the mathematical upper bound of variance explained by any collection of seven spatial topographies. Seven spatial components explained 95.03% (95% CI [94.59, 95.46]) of variance on average in the 128-channel EEG at the sensor level (see Supplementary Materials for depictions of the topographies). The effective spatial dimensionality was therefore seven components based on the 95% cumulative variance threshold. In contrast, seven PCA components explained 64.90% (95% CI [62.89, 66.92]) of variance on average directly among the distributed current densities at the source level, with the 95% threshold identifying an effective dimensionality of 44 source-level components. When the top seven sensor-level PCA components were subsequently source-localized, they explained 53.15% (95% CI [51.13, 55.18]) of the total source-level variance among the distributed current densities. But they accounted for 88.89% (95% CI [88.11, 89.67]) of the total variance in the reconstructed 128-channel EEG after source-level removal and forward-projection. The finding that seven microstate configurations account for roughly 83% and 91% of the variance in the 128-channel EEG on average demonstrates that the microstate configurations are near-optimal spatial representations for explaining the global signal relative to these limits. Notably, the spatial topography of the grand-average first principal component was nearly identical to the configuration of microstate C (spatial *r* = .998; see Supplementary Materials).

#### Removal of surrogate comparison topographies

Iteratively smoothed nearest-neighbor surrogate comparison topographies were generated to resemble EEG topographies that characterize genuine volume-conducted cortical dipoles. Removal of seven spatially smoothed noise surrogate topographies at the sensor level reduced 51.87% (95% CI [51.63, 52.11]) of variance on average across random permutations. In contrast, removing seven matched surrogate subspaces directly at the source level reduced only 16.31% (95% CI [15.96, 16.65]) of the variance among the distributed vector current densities within source space. After source-level removal and forward-projection of the topographies, the surrogates accounted for 8.65% (95% CI [8.34, 8.95]) of the total variance in the reconstructed 128-channel EEG. The variance explained by the surrogate topographies is much higher at the sensor level because the noise surrogates spatially overlap at random with physiologically meaningful activity and therefore capture a substantial share of variance and spectral power. However, source localization acts as a biophysically weighted spatial filter that heavily attenuates the influence of these artificial topographies. Source localization estimates them to have putative anatomical origins that are largely orthogonal to the modeled cortical dipoles estimated to underlie physiologically meaningful EEG. The fact that the seven microstate configurations explain nearly 83% of variance despite this transformation suggests they reflect modeled cortical dipoles that align with physiologically meaningful activity and explain a non-arbitrary portion of the EEG signal.

#### Power spectral density of the residual EEG

Next, we visually inspected the PSD of the original and residual EEG before and after removal of the modeled cortical sources of microstate configurations. Figure 4 depicts the PSD after source-level removal of configurations across both linear and semi-logarithmic scales for residual EEG compared to the original. The sensor-level PSD is provided in Supplementary Materials. Notably, the aggregate spatial filtering of the seven microstate configurations greatly attenuated the primary oscillatory peak in the alpha band (∼10 Hz) within the residual signal and resulted in global reduction in power across the entire broadband frequency range. Furthermore, visual inspection of the log-scaled spectra suggested an attenuation of the 1/*f* slope and reduction in the aperiodic exponent of the residual signal. Source-level spatial filtering of individual microstate configurations also revealed that specific configurations have variable contributions to the power spectrum. Microstate configuration C appeared to be a dominant contributor to both periodic and broadband power reductions. In contrast, removing microstates F and G had marginal effect on the spectra.

**Figure 4:**
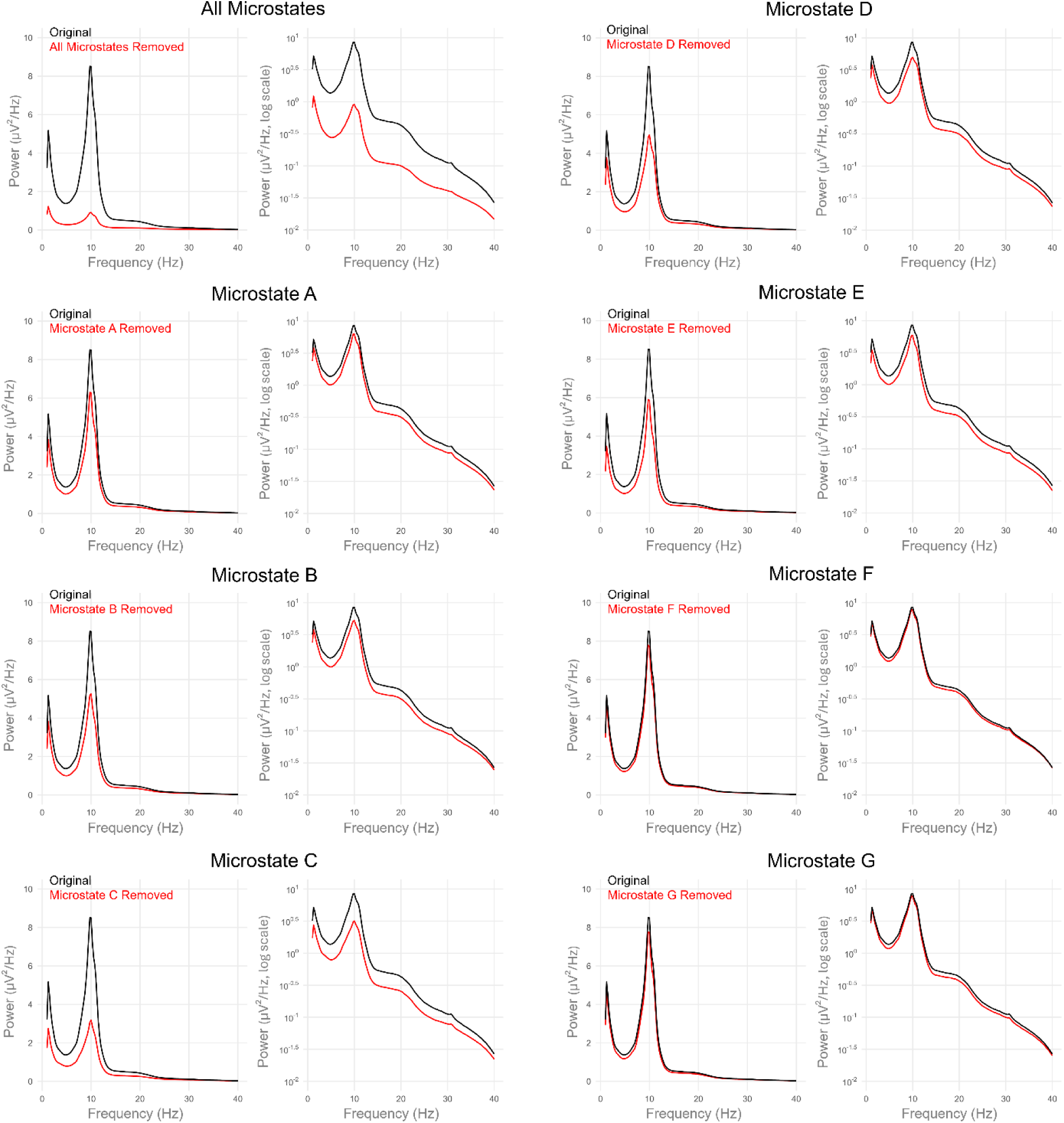
Power spectral density (PSD) of the original continuous EEG is shown compared to the residual signal following the computational removal of microstate topographies at the source level using orthogonal subspace projection (OSP). Each panel displays the PSD across the frequency range on both a linear scale (left) to emphasize the periodic oscillatory peaks and a semi-logarithmic scale (right) to emphasize the broadband aperiodic 1/*f* background. The intact EEG signal is depicted in black, while the residual signal is depicted in red following OSP of all seven microstates combined (top left) or the selective removal of individual microstate configurations A through G.

### Removing the Spatial Projections of Microstates Attenuates Parameterized EEG Spectra

Statistical comparisons of the parameterized power spectrum confirmed that both the periodic and aperiodic features of the macroscopic EEG depend on the spatial architecture of microstates.^3^ We first confirmed acceptable *specparam* model fit for the original EEG and after configurations were removed. The mean estimated *R*^2^ of parameterized models was 0.978 (95% CI [0.977, 0.980], range = 0.953 – 0.991) for the original data across recording blocks and 0.977 (95% CI [0.975, 0.978], range = 0.911 – 0.990) after source-level removal of all microstate configurations. The results of source-level removal of configurations are organized below according to the periodic and aperiodic spectral features, and the model-estimated effects are summarized in Table 2. Sensor-level removal of configurations broadly aligned with the source-level effects and are reported in Supplementary Materials. The reported results represent unstandardized beta coefficients comparing the residual EEG reconstructed after configurations were removed at the source level compared to the original EEG (i.e., the intercept). The threshold for reporting significant tests of parameter estimates was *p* < .001 to account for multiple comparisons across parameterized features of the residual EEG datasets.

**Table 2:** Model parameter estimates comparing periodic and aperiodic spectral parameters of the original to the residual EEG after source-level removal.

|  | Aperiodic-adjusted Alpha Power | Aperiodic-adjusted Theta Power | Aperiodic Offset | Aperiodic Exponent |
|---|---|---|---|---|
| Fixed Effects |  |  |  |  |
| Intercept (Original EEG) | 1.1295 (0.0431)*** | 0.4455 (0.0243)*** | 0.9500 (0.0346)*** | 1.4261 (0.0209)*** |
| All microstates removed | -0.3315 (0.0076)*** | -0.1268 (0.0053)*** | -0.8624 (0.0058)*** | -0.3185 (0.0050)*** |
| Microstate A removed | -0.0601 (0.0068)*** | -0.0245 (0.0049)*** | -0.1929 (0.0050)*** | -0.0689 (0.0045)*** |
| Microstate B removed | -0.0822 (0.0067)*** | -0.0295 (0.0048)*** | -0.2005 (0.0050)*** | -0.0812 (0.0045)*** |
| Microstate C removed | -0.1875 (0.0073)*** | -0.0572 (0.0049)*** | -0.3340 (0.0051)*** | -0.1368 (0.0046)*** |
| Microstate D removed | -0.0783 (0.0070)*** | -0.0374 (0.0050)*** | -0.2312 (0.0049)*** | -0.1038 (0.0045)*** |
| Microstate E removed | -0.0837 (0.0069)*** | -0.0278 (0.0049)*** | -0.2160 (0.0052)*** | -0.0650 (0.0046)*** |
| Microstate F removed | -0.0030 (0.0071) | -0.0160 (0.0053)** | -0.0917 (0.0051)*** | -0.0444 (0.0046)*** |
| Microstate G removed | 0.0066 (0.0072) | -0.0035 (0.0053) | -0.0932 (0.0051)*** | -0.0357 (0.0045)*** |
| Random Effects |  |  |  |  |
| Intercept variance | 0.1429 (0.0231) | 0.0444 (0.0072) | 0.0926 (0.0149) | 0.0334 (0.0054) |
| Residual variance<sub>Original</sub> | 0.0151 (0.0009) | 0.0062 (0.0004) | 0.0080 (0.0005) | 0.0062 (0.0004) |
| Residual variance<sub>All</sub> | 0.0210 (0.0013) | 0.0066 (0.0005) | 0.0129 (0.0008) | 0.0091 (0.0005) |
| Residual variance<sub>A</sub> | 0.0135 (0.0008) | 0.0054 (0.0003) | 0.0073 (0.0004) | 0.0061 (0.0003) |
| Residual variance<sub>B</sub> | 0.0128 (0.0007) | 0.0047 (0.0003) | 0.0076 (0.0004) | 0.0064 (0.0004) |
| Residual variance<sub>C</sub> | 0.0179 (0.0011) | 0.0052 (0.0003) | 0.0081 (0.0005) | 0.0070 (0.0004) |
| Residual variance<sub>D</sub> | 0.0151 (0.0009) | 0.0054 (0.0004) | 0.0071 (0.0004) | 0.0061 (0.0003) |
| Residual variance<sub>E</sub> | 0.0146 (0.0009) | 0.0053 (0.0003) | 0.0087 (0.0005) | 0.0068 (0.0004) |
| Residual variance<sub>F</sub> | 0.0165 (0.0010) | 0.0068 (0.0005) | 0.0080 (0.0005) | 0.0066 (0.0004) |
| Residual variance<sub>G</sub> | 0.0174 (0.0010) | 0.0069 (0.0005) | 0.0081 (0.0005) | 0.0066 (0.0004) |
| *N* | 78 | 77 | 78 | 78 |
| Obs. | 5580 | 4317 | 5580 | 5580 |

Note: Model parameters representing the fixed effects of removing specific microstate configurations from the original data using source-level orthogonal subspace projection on dependent measures of aperiodic-adjusted alpha power, theta power, aperiodic offset, and aperiodic exponent. Effects describe the difference between the residual signal and the original EEG (i.e., the intercept) for all microstates and each individual configuration A through G from multilevel mixed effects models. Random intercept variances and residual variances are also provided. Model estimates were based on a sample size of *N* = 78 with 5580 total observations contributing to analyses (*n* = 77 and 4317 observations for theta power). Standard errors are provided in parentheses. The statistical significance of fixed effects is indicated. \* *p* < .05, \*\* *p* < .01, \*\*\* *p* < .001

#### Periodic peak alpha and theta power

We observed a significant main effect of source-level removal of specific microstate configurations on aperiodic-adjusted alpha power, *F*(8, 2213) = 379.06, *p* < .001, and aperiodic-adjusted theta power, *F*(8, 1727) = 103.36, *p* < .001. The residual signal had significantly lower aperiodic-adjusted alpha power (*b* = −0.3315, *SE* = 0.0076, *p* < .001, 95% CI [-0.3465, −0.3166]) and theta power (*b* = −0.1268, *SE* = 0.0053, *p* < .001, 95% CI [-0.1372, −0.1164]) when all configurations were removed, compared to the original EEG (see Figure 5). Microstate configurations A, B, C, D, and E, reduced alpha power (*p*s < .001) when separately removed from the original EEG (see Table 2). In contrast, configurations F and G did not significantly change alpha power when removed. Similarly, all microstates except configurations F and G significantly (*ps* < .001) reduced theta power when removed from the original EEG (see Table 2). Removing configuration C reduced both alpha and theta power significantly (*p*s < .001) more than all other microstates. There were also variable contributions of specific microstates to the central frequency and bandwidth of the peak power in the alpha and theta bands (see Supplementary Materials). Overall, the targeted removal of microstate configurations revealed widespread reductions in both aperiodic-adjusted alpha and theta power for most configurations. There was a small subset of configurations (F and G) whose removal did not modulate these specific rhythms. Finally, the removal of microstate configuration C affected the periodic components of the PSD more than other microstates.

**Figure 5:**
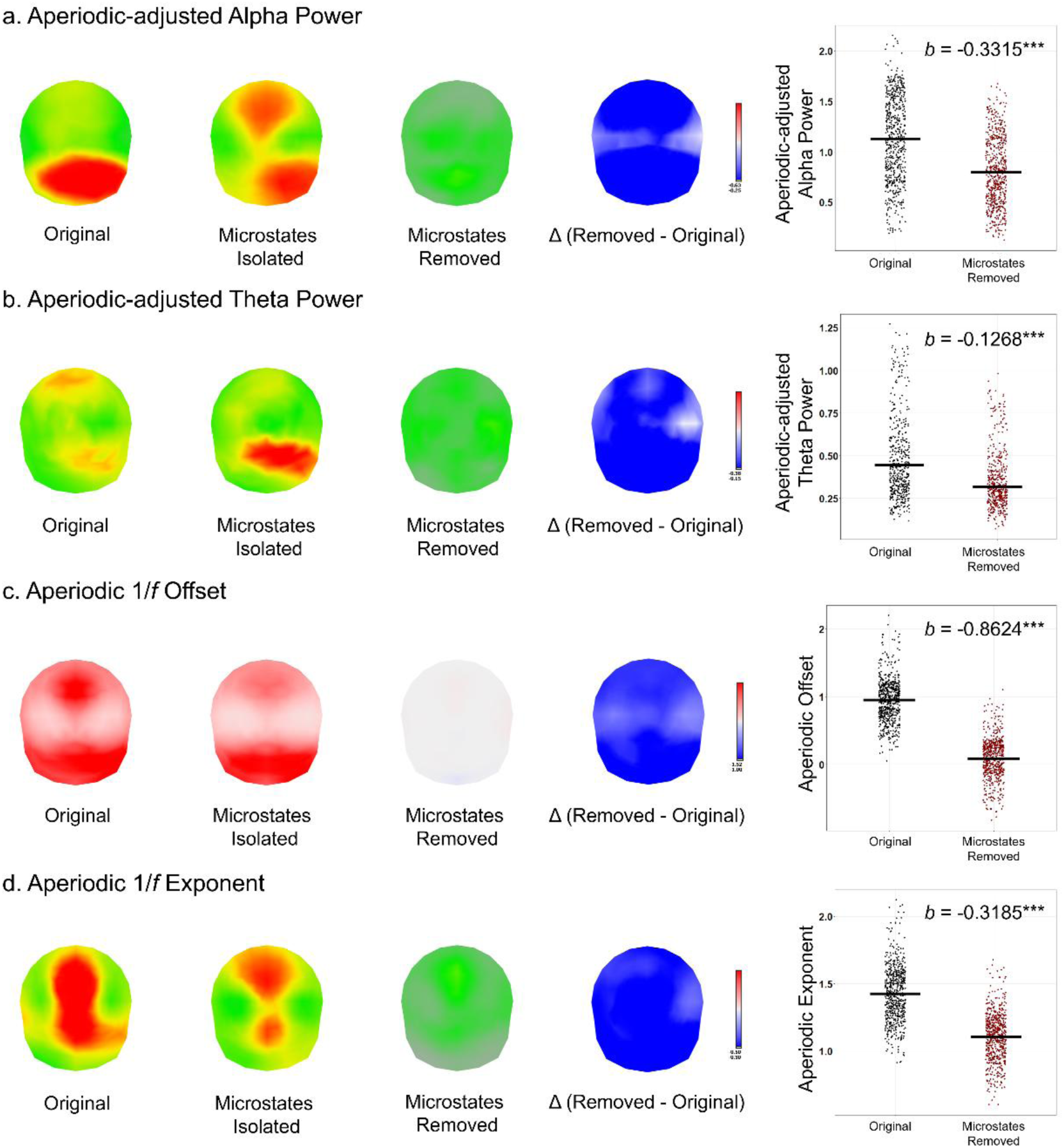
Topographic maps depict the grand average topography of the intact EEG signal (original), the grand average signal reconstructed from the source-level spatial projections of microstates (microstates isolated), the grand average residual signal after source-level computational removal of all microstates (microstates removed), and the model-estimated difference, Δ (removed – original), for a). the aperiodic-adjusted alpha power, b). the aperiodic-adjusted theta power, c). aperiodic 1/*f* offset, and d). aperiodic 1/*f* exponent. For the difference-maps, decreases are shown in blue at electrode locations and electrode locations with non-significant differences are omitted from the map. Jitter plots (right) display the corresponding individual data points from recording epochs for the original (black) and residual (red) signals averaged across the electrode montage. The horizontal black lines denote the model-estimated mean value. Unstandardized beta coefficients (*b*) denote the model-estimated reduction in each spectral parameter following the removal of all microstates. *** *p* < .001

#### Aperiodic 1/*f* offset and exponent

We observed a significant main effect of source-level removal of specific microstate configurations on the aperiodic offset, *F*(8, 2255) = 3400.40, *p* < .001, and aperiodic exponent, *F*(8, 2270) = 642.98, *p* < .001. The residual signal had significantly lower aperiodic offset (*b* = −0.8624, *SE* = 0.0058, *p* < .001, 95% CI [-0.8738, −0.8510]) and lower (flatter) aperiodic exponent (*b* = −0.3185, *SE* = 0.0050, *p* < .001, 95% CI [-0.3282, −0.3088]) when all configurations were removed, compared to the original EEG (see Figure 5). All microstate configurations reduced the aperiodic offset and aperiodic exponent when removed from the original EEG (see Table 2). Removing configuration C reduced the aperiodic offset and exponent significantly (*p*s < .001) more than all other microstates. Overall, the targeted removal of microstate configurations revealed widespread reductions in both the aperiodic offset and exponent for most configurations. The removal of microstate configuration C, however, affected the aperiodic components of the PSD more than other microstates.

#### Periodic and aperiodic spectral features of surrogate comparison topographies

We parameterized the PSD of the residual EEG following the removal of spatially smoothed surrogate noise topographies to confirm that the attenuation of spectral features was specific to the spatial architecture of microstates. The removal of surrogate topographies at the source level did not significantly alter the periodic peak alpha or theta power or the aperiodic exponent (see Table 3). There was only a small reduction in the broadband aperiodic offset, which is expected given the general reduction in signal variance (8.65%) associated with removing these topographies. This outcome is consistent with the inverse model effectively attenuating signals that do not overlap in source space with the actual physiological activity estimated to underlie these periodic and aperiodic components. On the other hand, the sensor-level surrogates ought to overlap at random with physiologically meaningful activity on the scalp and therefore capture some share of periodic and aperiodic spectral power. Removing the surrogates directly at the sensor level resulted in significant but comparatively small reductions to periodic alpha, theta, and aperiodic spectral features (see Supplementary Materials). However, these effects were substantially smaller in magnitude than those observed following the removal of the microstate configurations. Together, these control analyses demonstrate that the reductions in periodic rhythms and the flattening of the 1/*f* exponent observed in the primary analyses are specifically associated with the biologically plausible topographic configurations of EEG microstates.

**Table 3:**
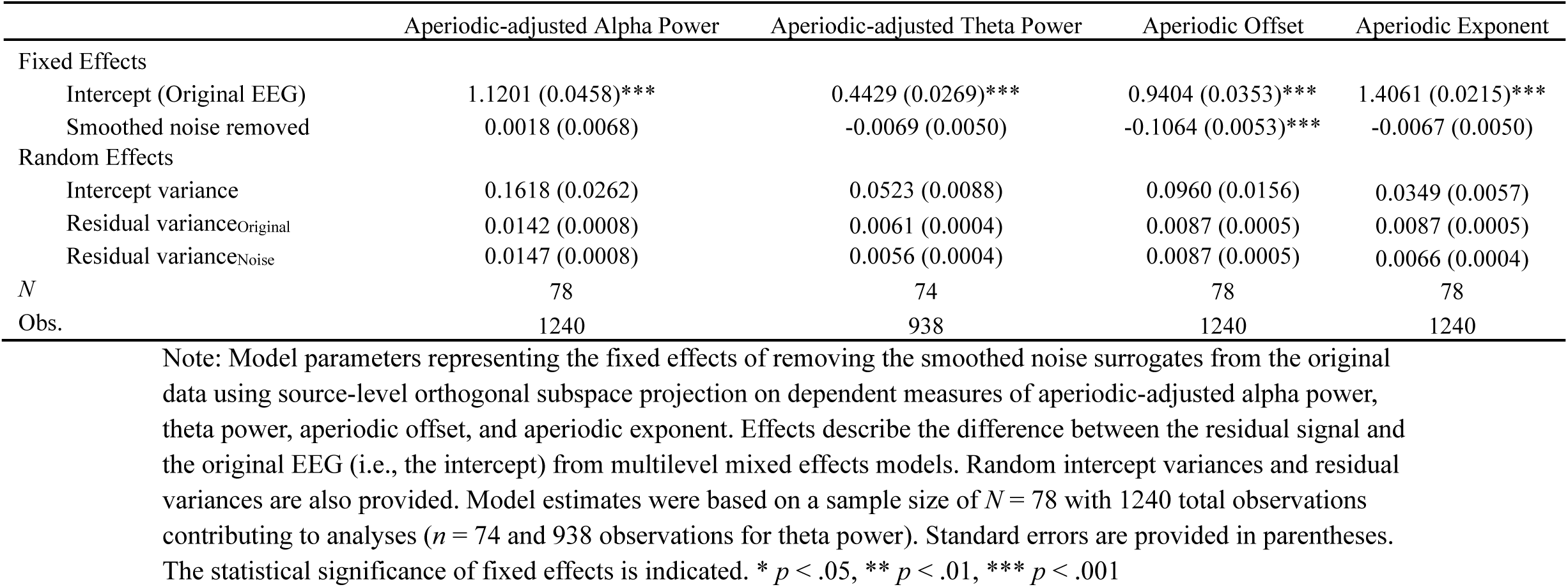
Model parameter estimates comparing spectral parameters of the original EEG to the smoothed noise surrogates after source-level removal.

### Removing the Spatial Projections of Microstates Attenuates Configuration Specific Spectra

Next, we evaluated the electrode-level effects of removing the spatial subspaces of microstates on the aperiodic-adjusted alpha power, aperiodic-adjusted theta power, aperiodic offset, and aperiodic exponent for all 128 electrodes. We therefore compared spectral estimates obtained for each electrode from the original continuous EEG time series to the same electrode following the targeted removal of subspaces of microstates at the source level through OSP. The results of sensor-level removal of microstates are generally consistent with source-level removal and are provided in Supplementary Materials. We observed that removing the spatial projections of all microstate configurations resulted in highly significant reductions in alpha power, theta power, aperiodic offset, and aperiodic exponent across most scalp electrodes (see Figure 5). Topographical representations of the original, isolated, residual, and difference signals are depicted in Figures 5 and 6. The isolated removal of individual microstate configurations, however, revealed spatially distributed effects. The removal of specific microstate configurations resulted in localized reductions in alpha and theta power, aperiodic offset, and aperiodic exponent, corresponding precisely to their unique spatial configurations (see Figure 6). In contrast to other configurations, the removal of microstate configurations F and G, which had minimal effect on overall mean alpha and theta power, demonstrated significant but spatially constrained differences in alpha power, theta power, aperiodic offset, and aperiodic exponent at a few electrode locations.

**Figure 6:**
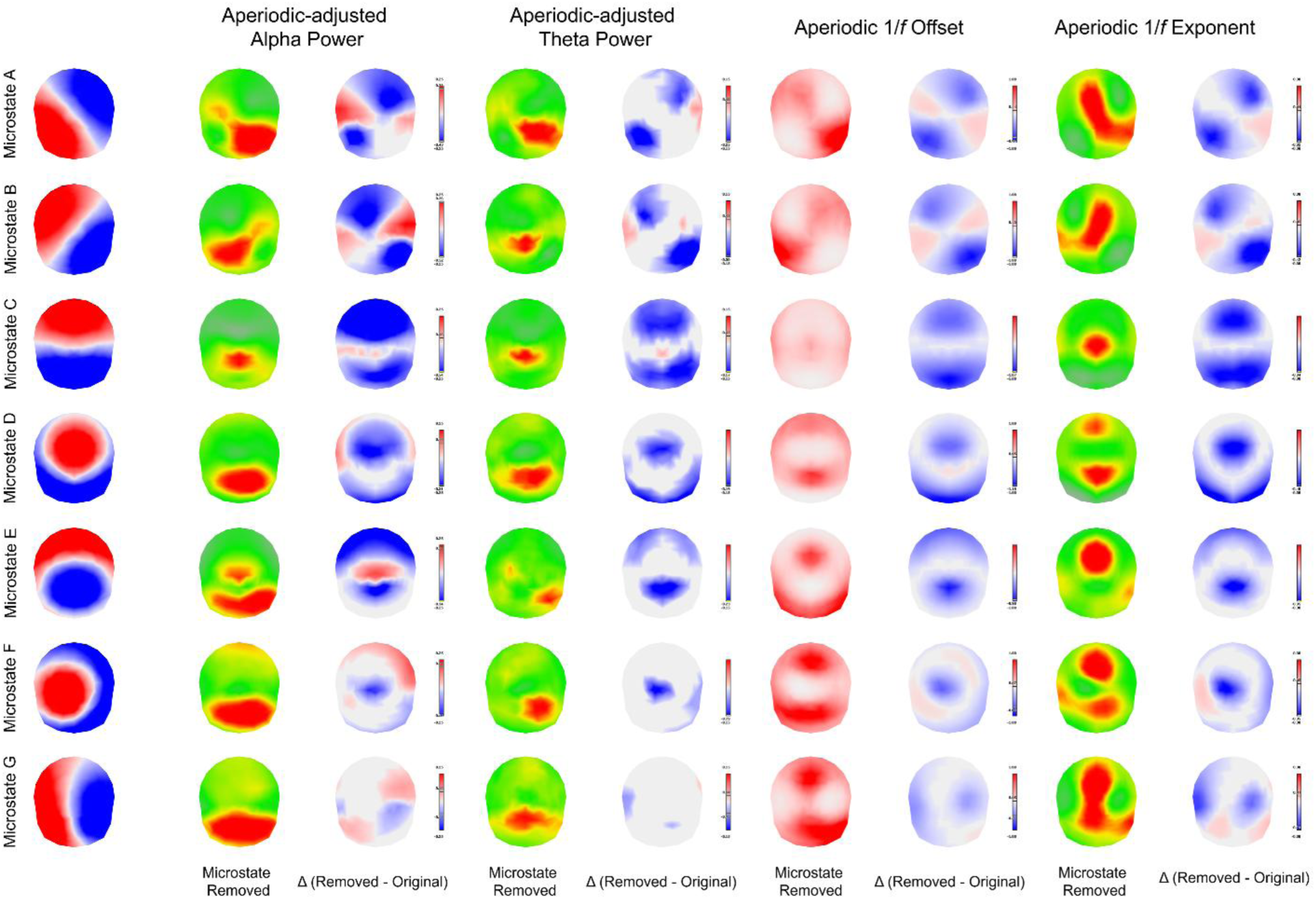
Topographic maps depict the spatially specific effects of computationally removing individual microstate configurations (A through G) at the source level. The leftmost column displays the voltage topography of each targeted microstate. The subsequent columns present the topography of grand average residual signal (microstate removed) and the model estimated difference, Δ (removed – original), for the aperiodic-adjusted alpha power, the aperiodic-adjusted theta power, aperiodic 1/*f* offset, and aperiodic 1/*f* exponent. Across all comparison Δ maps, significant increases are shown in red (indicating greater power at that electrode location when that microstate is removed), while decreases are shown in blue. Electrode locations with non-significant differences are omitted.

### Microstate Dynamics Predict Periodic and Aperiodic Spectral Features

We evaluated associations between spectral parameters and the lognormal distribution of microstate durations (lognormal µ and right-skewed shape σ) calculated both independently of polarity and as polarity-fixed segments to determine how the time-domain dynamics of microstates account for specific features of the EEG power spectrum. Model estimated means of microstate durations are provided in Table 4. Mixed-effects models identified several associations between microstate durations and periodic alpha power, theta power, and the broadband aperiodic offset and exponent. The simple effects are summarized in Table 5.

**Table 4:** Model estimated means for lognormal parameters of the microstate duration distribution.

| Microstate | Lognormal μ | Lognormal σ | Polarity-Fixed Lognormal μ | Polarity-Fixed Lognormal σ |
|---|---|---|---|---|
| All microstates | -2.6404 [-2.6494, -2.6313] | 0.6110 [0.6049, 0.6172] | -3.1924 [-3.2048, -3.1800] | 0.4163 [0.4059, 0.4268] |
| Microstate A | -2.7550 [-2.7645, -2.7455] | 0.5335 [0.5277, 0.5393] | -3.2429 [-3.2554, -3.2303] | 0.4062 [0.3957, 0.4167] |
| Microstate B | -2.7290 [-2.7384, -2.7196] | 0.5496 [0.5438, 0.5555] | -3.2360 [-3.2486, -3.2235] | 0.3974 [0.3869, 0.4079] |
| Microstate C | -2.4117 [-2.4241, -2.3994] | 0.6766 [0.6693, 0.6838] | -3.1605 [-3.1732, -3.1478] | 0.4300 [0.4192, 0.4407] |
| Microstate D | -2.7121 [-2.7220, -2.7022] | 0.5539 [0.5480, 0.5599] | -3.1877 [-3.2004, -3.1750] | 0.4161 [0.4055, 0.4266] |
| Microstate E | -2.7178 [-2.7280, -2.7075] | 0.5509 [0.5447, 0.5571] | -3.1826 [-3.1954, -3.1698] | 0.4248 [0.4142, 0.4354] |
| Microstate F | -2.8338 [-2.8433, -2.8243] | 0.4960 [0.4901, 0.5020] | -3.2076 [-3.2203, -3.1948] | 0.3883 [0.3777, 0.3988] |
| Microstate G | -2.8271 [-2.8365, -2.8177] | 0.4972 [0.4913, 0.5032] | -3.2507 [-3.2635, -3.2380] | 0.3813 [0.3707, 0.3918] |

Note: Model estimates of the mean lognormal location (μ) and shape (σ) of microstate durations (in seconds) are provided from multilevel mixed-effects models. Fixed effects of microstate configuration are estimated from 78 participants and 620 recording blocks of EEG. Durations were calculated ignoring polarity (lognormal μ and σ) and by accounting for spatial polarity (polarity-fixed lognormal μ and σ). 95% confidence intervals (CI) accompany the model estimated means.

**Table 5:**
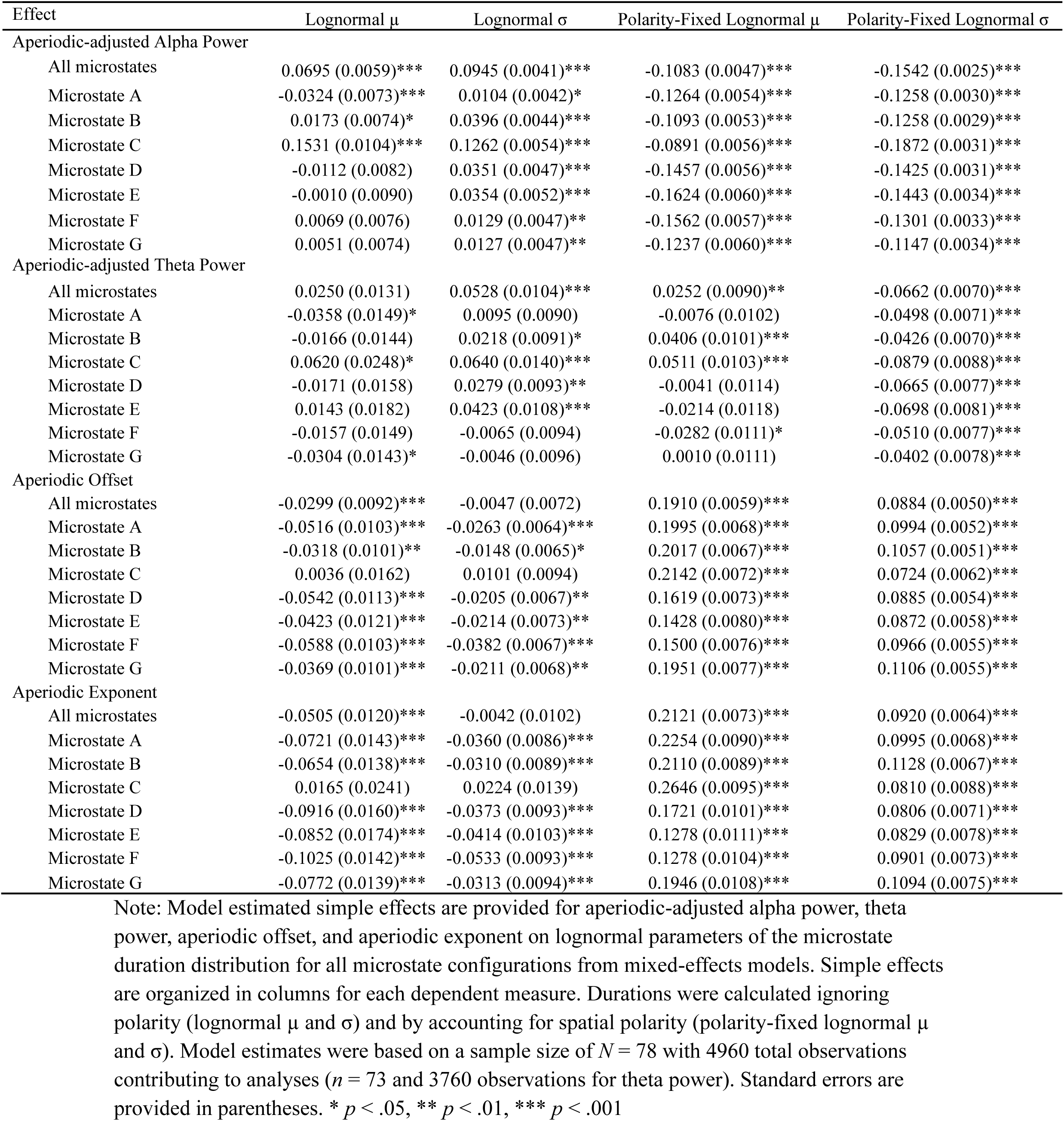
Simple effects of periodic alpha and theta power, aperiodic offset, and aperiodic exponent, predicting lognormal microstate durations.

Importantly, there were several noteworthy patterns confirming very strong associations between overall microstate durations and periodic and aperiodic spectral features (see Figure 7). First, overall polarity-independent durations of microstates were strongly associated with aperiodic-adjusted alpha power. Both the lognormal location µ (*b* = 0.0695, *SE* = 0.0059, *p* < .001, 95% CI [0.0578, 0.0811]) of durations and lognormal shape σ (*b* = 0.0945, *SE* = 0.0041, *p* < .001, 95% CI [0.0865, 0.1026]) were positively associated with alpha power, suggesting that microstates persist for long durations by oscillating in polarity within the alpha frequency range. In contrast, the lognormal shape σ of durations of polarity-fixed segments of microstates were negatively correlated with alpha power (*b* = −0.1542, *SE* = 0.0025, *p* < .001, 95% CI [-0.1591, −0.1493]) and theta power (*b* = −0.0662, *SE* = 0.0070, *p* < .001, 95% CI [-0.0799, −0.0526]). This suggests that increased variability in the durations of polarity-fixed microstate segments shift power out of periodic components into broadband frequencies. The lognormal µ duration of polarity-fixed segments of microstates was positively associated with greater aperiodic offset (*b* = 0.1910, *SE* = 0.0059, *p* < .001, 95% CI [0.1795, 0.2026]) and steeper aperiodic exponent (*b* = 0.2121, *SE* = 0.0073, *p* < .001, 95% CI [0.1979, 0.2264]).

**Figure 7:**
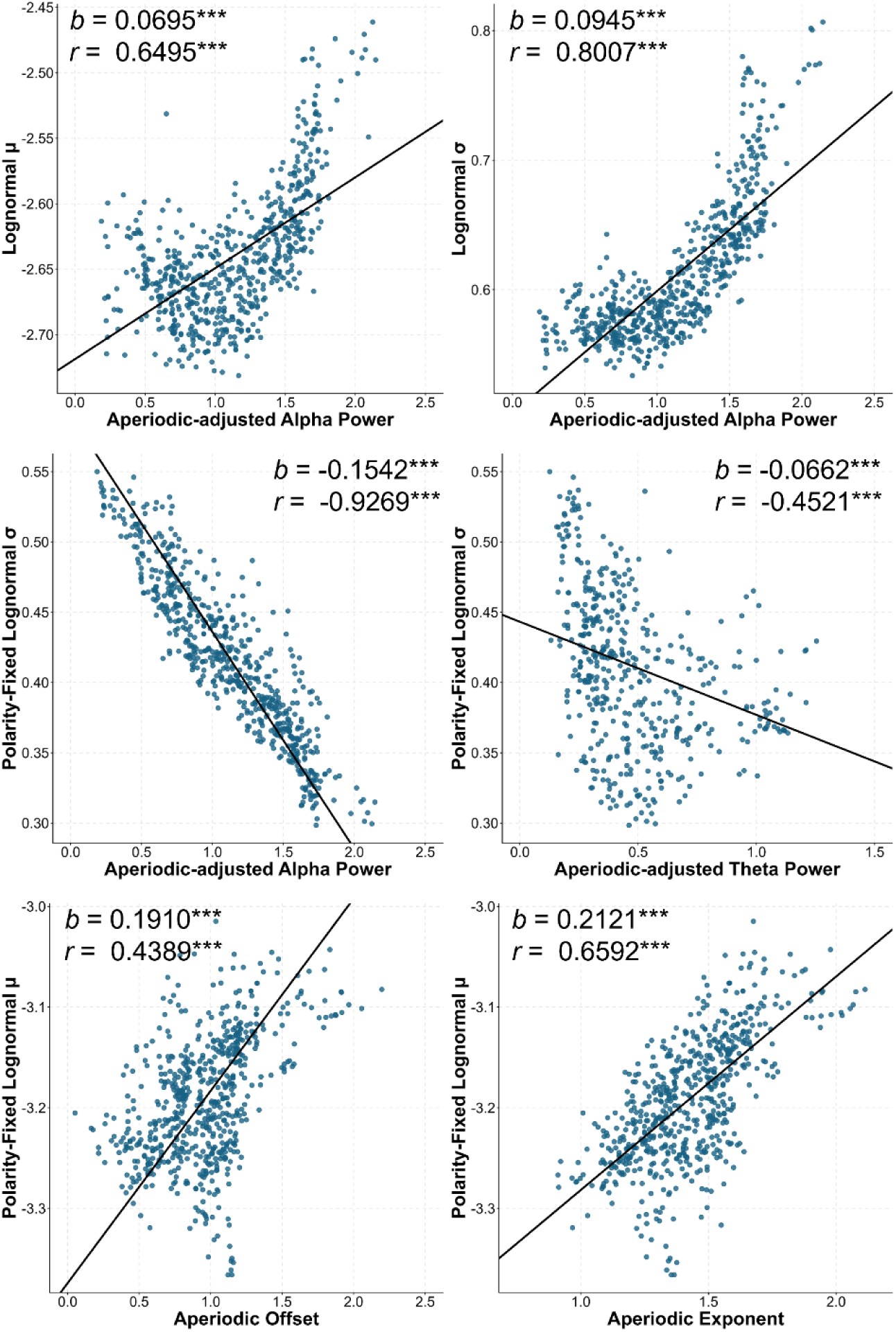
Scatterplots depict associations from mixed-effects models between the lognormal location (µ) and shape (σ) of the microstate duration distribution and periodic and aperiodic spectral parameters. Lognormal microstate durations were calculated ignoring polarity (lognormal µ and σ) and by accounting for polarity (polarity-fixed lognormal µ and σ). Model estimated intercepts and slopes describing associations between variables are plotted on top of the observed data. The slope (*b*) estimate is provided. The between-subject correlation (*r*) is provided from values averaged across recording blocks. *** *p* < .001

To contextualize the magnitude of these effects, we averaged values across recording blocks for individuals and computed between-subject correlations. At the subject-average level, lognormal µ duration (*r* = 0.6495, *p* < .001, 95% CI [0.4991, 0.7619]) and lognormal σ (*r* = 0.8007, *p* < .001, 95% CI [0.7035, 0.8685]) had strong positive correlations with mean aperiodic-adjusted alpha power. The associations between polarity-independent microstate durations and aperiodic-adjusted alpha power appeared to follow a nonlinear quadratic pattern (see Figure 7). Adding a quadratic model term explained 66.62% of the total variance (*R*^2^ = .6662, *R* = 0.8162, *p* < .001) for lognormal µ duration and 78.78% (*R*^2^ = .7878, *R* = 0.8876, *p* < .001) for lognormal σ. This suggests that individuals with greater levels of periodic alpha power are characterized by microstates that increasingly persist for long durations by oscillating in polarity in the alpha frequency range. The polarity-fixed lognormal σ, however, had strong negative correlations with mean alpha power (*r* = −0.9269, *p* < .001, 95% CI [-0.9529, −0.8874]) and theta power (*r* = −0.4521, *p* < .001, 95% CI [-0.6179, −0.2478]), explaining 85.91% and 20.44% of between-person variance, respectively. Finally, polarity-fixed lognormal µ had a strong positive correlation with aperiodic offset (*r* = 0.4389, *p* < .001, 95% CI [0.2398, 0.6026]) and exponent (*r* = 0.6592, *p* < .001, 95% CI [0.5117, 0.7689]), explaining 19.27% and 43.44% of between-person variance, respectively.

Examining the associations of individual microstate configurations revealed their specific contributions to these effects (see Table 5). For polarity-fixed measures of duration, most microstates were similarly associated with spectral features. One exception was associations of polarity-independent durations of microstate C with aperiodic-adjusted alpha power. These associations were significantly more positive than other microstates (*p*s < .001), suggesting that long-duration microstates associated with activity in the alpha band are strongly influenced by instances of microstate C persisting in time by successively oscillating in polarity. Microstate dynamics and periodic and aperiodic spectral features therefore demonstrate a high degree of statistical redundancy. This supports our explanatory model for how time-domain dynamics of microstates structurally constitute the periodic and aperiodic features of the neural power spectrum.

## Discussion

Isolating and computationally removing the specific source- and sensor-level spatial subspaces occupied by seven microstate configurations revealed that the states represented by these configurations contribute to both periodic and aperiodic features of the EEG neural power spectrum. Notably, removing all seven configurations together greatly attenuated the alpha power peak and reduced the aperiodic 1/*f* offset and exponent. This suggests that the time-domain behavior of the microstates associated with these spatial configurations constitutes most of the periodic and aperiodic features of the broadband EEG frequency spectrum. The seven microstate configurations explained roughly 83% and 91% of the total signal variance when removed from the 128-channel eyes-closed resting EEG through source- and sensor-level OSP, respectively, demonstrating that these topographic configurations are near-optimal spatial representations explaining global signal variance relative to the theoretical upper limit established by PCA. This provides compelling evidence that the repertoire of configurations encompasses a substantial portion of the brain’s broadband and alpha-generating macroscale network architecture. In this way, our findings offer a reimagining of Lehmann’s (1990) original vision of microstates as the “atoms of thought” by positioning them as a fundamental spatial basis set describing the majority of structured periodic and aperiodic dynamics observable in the scalp electric field.

The contributions of specific microstates to the spectral content were not uniform. We found that the removal of most microstate configurations attenuated alpha and theta rhythms in the resulting residual signal. The removal of microstate configuration C, however, led to a greater proportional reduction in variance than other microstates, and reduced aperiodic-adjusted alpha and theta power to a greater extent. Its removal also attenuated the broadband aperiodic offset and exponent to a greater degree than other microstates, suggesting that this microstate contributes a steeper aperiodic 1/*f* exponent and more low frequency power to the spectrum than others. Independent corroboration through cross-spectral modeling also demonstrates that microstate C is one of the strongest contributors to the broadband EEG cross-spectrum, with its peak contribution occurring in the alpha band (Pascual-Marqui et al., 2022).

This is consistent with the predominance of this microstate within the EEG time series: microstate configuration C occupies the largest share of the EEG time series, occurs most frequently, and has the longest average duration (see Table 4; Zanesco, 2024). In fact, the topographic configuration of microstate C was nearly identical (spatial *r* = .998) to the topography of the grand average first principal component derived from the mean spatial covariance matrix across all subjects. This demonstrates that microstate configuration C is functionally equivalent to the principal component representing the primary spatial axis of EEG signal variance. The vast majority of signal variance is therefore anchored around one specific, recurring configuration. This is reminiscent of neuroimaging observations demonstrating that the first principal functional connectivity gradient defines a cortical axis separating transmodal association areas, anchored around the default mode network, from sensory-motor regions (Huntenburg et al., 2018; Margulies et al., 2016). This parallel implies an intriguing hypothesis that microstate C may represent an electrophysiological analog of this primary functional axis.

Projecting the spatial configurations of microstates F and G out of the EEG, however, did not significantly reduce alpha or theta on average across the electrode montage and instead reduced only the overall broadband aperiodic offset and exponent. This is consistent with the fact that these microstates explain significantly less topographic variance than others and are frequently excluded from the most common four- or five-configuration models of microstates (Koenig et al., 2024; Michel & Koenig, 2018). While removal of microstates F and G did not significantly reduce average alpha or theta power, individual differences in their durations were nevertheless associated with between-person variance in power estimates for these periodic rhythms. Importantly, removal of these configurations also serves as a methodological validation demonstrating the spatially selective removal of variance using OSP. Specifically, spatial filtering isolated the electrophysiological activity unique to each microstate configuration rather than indiscriminately suppressing the entire voltage recording and resulting spectra.

Microstate dynamics are expected to contribute to the neural power spectrum given that they predominate in the time-domain signal and their behavior manifests both as transient occurrences and as sustained oscillations. Microstates often persist in time because they oscillate in the alpha frequency range by inverting in polarity for several oscillatory cycles. These sustained oscillations are described as temporally extended, polarity-independent occurrences by standard microstate categorization approaches. The prevalence of temporally extended microstates was strongly correlated with alpha band power, suggesting that long-duration microstates are sustained by oscillatory behavior in the alpha band. Both the mean log-duration and the dispersion of the duration distribution’s long right tail were positively associated with aperiodic-adjusted alpha power. In contrast, when microstates are analyzed as fixed-polarity segments, spectral power is distributed out of the alpha band and into slower frequencies when half-cycle segments are longer in duration. The dispersion of the polarity-fixed distribution was therefore strongly negatively correlated with aperiodic-adjusted alpha power, whereas the mean log-duration of fixed-polarity segments was positively associated with aperiodic offsets and exponents across individuals. Roughly 86% of between-person differences in alpha power are explained by the average temporal persistence of their microstates. This shared variance indicates that time-domain measures of microstate durations and periodic alpha power are psychometrically redundant measurements of the same underlying brain dynamics (Rönkkö & Cho, 2022).

The time-varying dynamics of microstates provide temporal mechanisms linking their behaviors to spectral power. Microstate durations explain the vast majority of variance in periodic alpha power, but these measures account for a more modest share of between-person variance (roughly 43%) in the aperiodic exponent. We speculate the remaining variance may be explained by the temporal envelopes of time-varying oscillatory bursts and the temporal autocorrelation and long-range memory of microstate sequences. Specifically, the 1/*f* aperiodic exponent is the mathematical equivalent to long-range temporal correlations present in the time-domain. The scaling behavior of microstate sequences, quantified through measures such as the Hurst exponent, is a highly probable mechanism underlying aperiodic spectral features. Information-theoretic approaches have revealed that microstate sequences possess extended temporal dependencies that serve as plausible time-domain constituents of these spectral features (von Wegner et al., 2017, 2021). Likewise, quantifying the duration and variability of oscillatory bursts of microstates may explain some meaningful proportion of variance in the aperiodic exponent. Future work will therefore need to integrate approaches for characterizing these dynamics and temporal sequence dependencies to fully explain the temporal mechanism connecting microstates to the aperiodic offset and exponent. Sequential transitions between microstates have also been associated with other oscillatory phenomena, such as rotating phase patterns and the propagation of traveling waves (Li et al., 2025; von Wegner et al., 2021), providing some indication that the dynamics of microstates might be associated with traveling waves and non-zero-lag phase connectivity in the alpha band.

### Implications of Microstates as Functional Constituents of the EEG Power Spectrum

The spatial distribution of average aperiodic-adjusted alpha and theta power within the residual signal recapitulates the voltage topographies of each microstate configuration (see also Zanesco, Gross, et al., 2026). Power associated with microstate configuration C, for instance, overlapped with anterior and posterior electrode locations and its removal at both the source and sensor level heavily attenuated posterior alpha power and signs of the canonical posterior alpha rhythm. Importantly, this configuration was not the sole mediator of posterior alpha. Several other microstates were also associated with alpha power at posterior scalp locations. This demonstrates the state-mixing problem inherent to traditional spectral analyses. Time-aggregate estimates of alpha power, theta power, and components of the 1/*f* aperiodic background reflect multivariate mixtures of neural events occurring in the time-domain, including functionally distinct microstates. This consideration is an extension of the so-called “Fourier fallacy” (Jasper, 1948; Preston et al., 2026): spectral decomposition alone cannot differentiate the functionally distinct neural events that constitute those spectral estimates. Instead, the real-time fluctuations of the brain’s electric fields hold ontological priority, and spectral estimates are only mathematical descriptions used to decompose and summarize those dynamics.

Theoretical models positing *alpha oscillations* as a singular brain mechanism responsible for global cortical idling or sensory gating (e.g., Foxe & Snyder, 2011; Jensen & Mazaheri, 2010; Klimesch, 2012; Klimesch et al., 2007) must be reevaluated to account for the functionally distinct events that underlie those rhythms in EEG. The established relevance of microstates for human cognition would imply that alpha oscillations serve varying cognitive functions because they are inherently tied to the behavior of distinct brain circuits generating those rhythms at any given moment. It is possible that differences in the prevalence and dynamics of specific microstates mediate a broad range of psychologically relevant spectral phenomena. We hypothesize that diverse spectral effects, including increased alpha power during the maintenance of information in working memory (e.g., Jensen et al., 2002), as well as associations of frontocentral theta with attention and cognition (e.g., Cavanagh & Frank, 2014), and broadband aperiodic components associated with clinical and neurodevelopmental disorders (e.g., Karalunas et al., 2022; Ostlund et al., 2021), can be explained by changes in the dynamics of one or more microstates.

Increased alpha power over posterior scalp locations has been alternatively associated with successful maintenance of representations in working memory (Bonnefond & Jensen, 2012; Jensen et al., 2002; Roux & Uhlhaas, 2014), as well as attentional and perceptual failures and the occurrence of task-unrelated thought (Kam et al., 2022; Samaha et al., 2020; Van Diepen et al., 2019). These seemingly contradictory associations might be reconciled by identifying the distinct microstates that predominate in each context. In recent work, for example, we demonstrated that the competing dynamics of microstates C and E contribute to states of inattention and focus, respectively, and likely mediate attention-related differences in prestimulus alpha power and oscillations in tasks requiring sustained attention (Zanesco, Gross, et al., 2026). Alpha power, alpha oscillations, and prevalence of microstate C were all greater in the moments preceding stimulus detection errors and self-reported inattention. This alignment is unsurprising given that the topography of microstate C correlates with the first spatial principal component of the EEG signal and contributes heavily to posterior alpha power.

Other spectral characteristics of EEG that are associated with cognition or signatures of specific states of consciousness, such as sleep-related slow waves, may also originate within the brain network architecture defined by a subset of microstate topographic configurations. Indeed, microstates have been shown to predominate and explain a substantial portion of topographic variance during sleep (Brodbeck et al., 2012; Wiemers et al., 2024) and anaesthetized states (Artoni et al., 2022; Hermann et al., 2024). EEG microstates lengthen in duration on average during sleep with periodicities coupled to specific EEG frequencies across sleep stages (Wiemers et al., 2024). Other research has suggested that changes in the predominance of specific microstates may explain the characteristic anteriorization of alpha power during propofol-induced anesthesia (Shi et al., 2020). Together, these findings imply that sleep-related slow waves and other spectral signatures of altered states are intrinsically anchored within the large-scale network architecture underlying EEG microstates. The computational approach employed here provides one possible method to directly test whether the selective removal of microstates attenuates state-specific spectral signatures.

### Methodological Validation, Interpretation, and Limitations

We acknowledge that our explanatory framework applies to intrinsic time-varying EEG dynamics but that event-related activity, including stimulus-evoked modulations in power and phase-locking, may operate under different mechanistic principles. Furthermore, our framework is an organizational model of the scalp electric field and does not explain the precise thalamocortical mechanisms, cellular processes, or neuromodulatory influences that generate and sustain distinct microstates. Principles governing the intracortical rhythms of regionally specific local field potentials do not necessarily apply to the globally synchronized activity responsible for large-amplitude electric field configurations observable at the scalp. The EEG signal is heavily influenced by the activity of global cortical networks, whereas most of the brain’s neuronal activity is thought to be fundamentally invisible to scalp recordings due to closed-field generator geometries. As such, our claims are constrained to the macroscopic features of the dominant frequency spectrum (< 30 Hz) in human EEG. It is known that faster, more localized rhythms, such as specific beta-band generators and activity in the gamma band, operate under different biophysical constraints than the slower fluctuations that dominate the scalp EEG time series (Buzsáki & Draguhn, 2004; Buzsáki et al., 2013; von Stein & Sarnthein, 2000). Higher-frequency fluctuations are too low amplitude to strongly determine the global topographic pattern of the scalp electric field to the same degree as slower, larger-amplitude fluctuations.

Neurophysiological perspectives identify the slope of the 1/*f* broadband curve as a putative marker of asynchronous baseline neuronal fluctuations and the excitation-inhibition balance within cortical networks (Gao et al., 2017; Preston et al., 2026). Transient neural events and time-varying oscillatory bursts are frequently considered signal confounds from this perspective because they also contribute broadband power that obscures estimation of “true” aperiodic activity (Brake et al., 2024; Gyurkovics et al., 2022; Preston et al., 2026). While we remain agnostic as to the cellular and synaptic mechanisms underlying aperiodic activity, our findings provide empirical support for alternative models proposing that periodic and aperiodic features of the EEG power spectrum primarily result from the dynamics of slow oscillations and propagating arrhythmic events arising from the coordinated activity of large-scale brain networks (Brake et al., 2024; Buzsáki & Draguhn, 2004; Buzsáki & Vöröslakos, 2023). Indeed, we argue that microstate dynamics are associated with the aperiodic offset and exponent precisely because of their transient, irregular occurrences and time-varying oscillatory bursts. Future research might investigate the extent to which microstates and the aperiodic background are associated independent of these macroscale dynamics.

Orthogonal subspace projection was used in an attempt to isolate the spectral contributions of distinct microstates. We applied the projections at both the source and sensor levels because the continuous scalp electric field is a mixture of volume conducted sources that make sensor-level spatial filtering prone to unintended signal loss. This occurs because volume conduction from distributed brain sources to the same surface electrodes can cause unrelated neural activity to be attenuated if it aligns with the spatial weights of the targeted microstate configuration. Indeed, while the sensor-level analyses (see Supplementary Materials) aligned with the patterns identified using source-level OSP, the residual signal suffered from greater overall attenuation. Source-level OSP was an attempt to mitigate unintended signal loss caused by removing spatially correlated topographies by instead manipulating the specific 3D dipole orientations and current density magnitudes associated with the microstate configurations within the modeled cortical volume (Michel & Brunet, 2019; Schoffelen & Gross, 2009). This allowed activity from other neural processes within the same modeled voxels to remain unaltered if the estimated 3D dipole orientation was geometrically orthogonal to the spatial subspace of the targeted microstate configuration. It is therefore significantly less likely that unrelated neural signals are removed due to spatial overlap occurring from volume conduction at the scalp.

It is important to emphasize, however, that we utilized the inverse solution to conduct OSP at the source level primarily as a biophysically-weighted spatial filter rather than in an attempt to identify the true anatomical components of the neural generators of microstates. Source localization only provides a modeled approximation of neural activity rather than definitive anatomical solutions. However, we must acknowledge that real cortical functional networks are known to share overlapping neuronal populations and anatomical nodes. Furthermore, the sensor-to-source transition relies on mathematical constraints and biological assumptions regarding source depth and spatial smoothness, which also introduce source leakage (Hauk et al., 2011; Palva et al., 2018). The consequence is that projecting out the variance associated with one microstate at the source level may still inadvertently capture and remove variance from other concurrent neural processes. Thus, estimates of configuration-specific variance are imprecise. Nevertheless, biophysical laws dictate microstate configurations must be generated by at least partially distinct neural networks in order for their scalp topographies to differ (Vaughan, 1982).

Spatial leakage at the sensor level was explicitly demonstrated through our analyses of randomly generated smoothed noise surrogate topographies. Noise surrogates spatially overlap at random with topographies generated by physiologically meaningful activity and therefore capture some share of variance and spectral power. Signal variance was reduced by nearly 52% at the sensor level when these topographies were removed with OSP. Source-level removal, however, mitigated the amount of variance explained by surrogates. Forward projecting these topographies after their source-level removal explained only 9% of variance at the sensor level. This demonstrates that the noise surrogates largely fail to survive the sensor-to-source transformation because they do not correspond with plausible cortical generator configurations within the leadfield, and thus minimally influence the reconstructed EEG. In contrast, signal projection of the microstate configurations still accounted for 83% of variance when the EEG was reconstructed after source-level removal. Microstate configurations survive the sensor-to-source transformation because their putative source estimates reflect cortical activity that aligns both with the biophysical constraints of the head model and the actual empirical activity underlying the EEG.

OSP projections used a set of seven group-level topographic microstate configurations derived from sparse sampling of GFP peaks within individual recordings. The topographies closely replicate normative meta-analytic configurations established across independent laboratories (Koenig et al., 2024). As such, they represent generalizable topographic configurations that are not individually optimized to explain idiosyncratic subject-specific variance. Fundamentally, the microstate model is a time-domain approach intended to identify real physiological events occurring in the EEG (Michel & Koenig, 2018; Michel & Bréchet, 2026). This constrasts with blind source separation methods that are optimized to maximize explained statistical variance and derive components regardless of their biological plausibility. That a fixed, group-invariant topographic basis set of microstate configurations was able to explain near-ceiling signal variance across individual recordings supports both the validity of the microstate model and the notion that their spectral contributions reflect genuine biologically structured neural dynamics.

## Conclusions

Frequency-domain analyses remain a highly valuable and complementary approach to understanding the global signal characteristics of brain electrical activity. Recent frameworks parameterizing the EEG power spectrum into periodic oscillations and aperiodic components describe distinct characteristics of global brain activity that may not be immediately apparent in the dynamics of microstates (Donoghue et al., 2020; Ostlund et al., 2022). Shifts in spectral power and broadband aperiodic parameters undoubtedly provide useful insights into the brain’s global dynamic tendencies, neurochemical balance, and structural network constraints (Gao et al., 2017; Preston et al., 2026). Nevertheless, our findings reframe the interpretation of time-averaged EEG spectral estimates and challenge researchers to carefully consider the time-domain events that constitute the aggregate signal. The strength of microstate analysis is its ability to isolate state-specific patterns of global brain activity directly in the time domain. Time-averaged spectral estimates that serve as the primary biomarkers of human EEG necessarily conflate the spectral contributions of distinct microstates. Spectral methods therefore assess the aggregate influence of these states resulting from their global average dynamics.

In line with compelling correlational (e.g., Croce et al., 2020; Dziego et al., 2024; Javed et al., 2019; Milz et al., 2017; von Wegner et al., 2017, 2021; Zanesco, Gross, et al., 2026) and mathematical (Pascual-Marqui et al., 2022) evidence, we demonstrate that the time-domain dynamics of microstates constitute the periodic and aperiodic features of the broadband EEG frequency spectrum. While microstates account for most of the periodic alpha power, their different behavioral regimes also contribute power throughout broadband frequencies. Oscillations in the alpha range describe just one aspect of their complex temporal behavior. Although the importance of microstates as time-domain constituents of the EEG power spectrum has long been implicitly understood within the field of research on microstates (Lehmann et al., 1987), the implications of this perspective warrant broader recognition and acceptance. In the future, applications relying on spectral biomarkers must account for the fact that frequency-domain features are strongly associated with the temporal dynamics of microstates. Simultaneously, acknowledging the connection between EEG spectra and microstates provides new directions for investigating their functional significance by leveraging the extensive research on the cognitive, clinical, and developmental correlates of EEG spectral features.

## Acknowledgements

We thank Delanie Spivey, Abigail Gross, Lexi Horn, Braxton Stevenson, and Sophia Zanelli for their help collecting these data. We utilized the freely available Cartool EEG software toolbox (cartool.unige.ch) developed by Denis Brunet at the Functional Brain Mapping Lab, then at the Epilepsy and Networks Lab, University of Geneva, Switzerland, and supported by the Center for Biomedical Imaging (CIBM), Switzerland.

## Author Contributions

Anthony P. Zanesco: Conceptualization, Data curation, Formal analysis, Funding acquisition, Investigation, Methodology, Project administration, Resources, Visualization, Writing - original draft. Rene A. Perez: Conceptualization, Writing - review & editing.

## Data and Code Availability

The data that support the findings of this study are available in the OSF repository and can be found at: https://osf.io/qz859. EEG processing and source localization were performed using the freely available Cartool software toolbox (cartool.unige.ch).

## Ethics

The study was approved by the Institutional Review Board of the University of Kentucky. All participants provided written informed consent prior to participating in the study.

## Footnotes

1 The microstate configurations from the seven clusters could be identified among the meta-analytic topographies reported by Koenig et al. (2024) based on their strongest spatial correlation (*r* range = .932 – .992). One exception was microstate configuration F which only moderately correlated (*r* = .653) with one of the meta-analytic topographies. This indicates that configuration F was likely more idiosyncratic to our study. Nevertheless, we chose to retain the seven-cluster solution because similar lateralized topographies have been reported in the broader literature (e.g., Custo et al., 2017; Zanesco, Skwara, et al., 2021).

2 The proportional reduction in variance differs mathematically from topographic global explained variance (GEV) that is commonly used to quantify the explained variance of microstates (Murray et al., 2008). GEV is calculated from a mutually exclusive “winner-takes-all” categorization of scalp voltage topography based on each GFP-normalized sample’s spatial correlation with the global topographic configuration of each microstate.

3 Spectral parameterization of the PSD identified periodic peak components within the alpha band in 125.67 (*SD* = 10.13) electrodes on average across all recording blocks from the original and residual EEG. Although not visible in the grand average PSD, peaks in the theta band were identified in 45.47 (*SD* = 39.44) electrodes on average across all recording blocks from the original and residual EEG. One individual was excluded from analyses of theta (and four from the surrogate theta comparisons) because none of their recording blocks contained enough identifiable theta peaks across all conditions. Five individuals did not have enough identifiable theta peaks when their original EEG was considered alone. Aperiodic components were detected in all 128 electrodes in the recording blocks of every condition.

