## Supplemental Materials for "Macroscale dynamics of EEG microstates constitute the periodic and aperiodic features of the neural power spectrum"

### Supplementary Materials

#### Lower-Order Microstate Clustering Solutions

Topographic clustering of microstates led us to select the seven-cluster solution. We also examined the microstate configurations of the four-, five-, and six-cluster solutions to evaluate whether lower order subsets of configurations were contained within the seven-cluster solution. The microstate configurations from the four-cluster solution (spatial *r*s > .942), five-cluster solution (*r*s > .992), and six-cluster solution (*r*s > .977), were all strongly spatially correlated with specific configurations in the seven-cluster solution. There was one exception in the six-cluster solution. The sixth configuration showed a somewhat lower correlation with configuration F ($r = .776$) prior to the separation of configurations F and G in the seven-cluster solution. Supplementary Figure 1 depicts the topographic configurations of microstates from the seven-cluster microstate solution and lower-order cluster solutions. This indicates that subspace projections using lower *k* cluster solutions would yield nearly identical configuration-specific results because the lower cluster solutions are fully encompassed by configurations in the seven-cluster solution.

#### Principal Component Topographies

Principal component analysis (PCA) confirmed the spatial dimensionality of the sensor and source EEG. The eigenvectors representing the spatial topographies of the first seven components were extracted from the 128-channel recordings and visualized using the mean spatial covariance matrix across all individuals. The seven PCA topographies are shown in Supplementary Figure 2. Notably, the spatial topography of the grand-average first principal component was nearly identical to the configuration of microstate C (spatial $r = .998$).

**Spatially Smoothed Surrogate Topographies**

A surrogate analysis was conducted using topographies generated from spatially smoothed noise. A total of 10,000 independent topographies were randomly generated based on values sampled from a standard Gaussian distribution ($\mu = 0$, $\sigma = 1$). The values were iteratively smoothed until topographies resembled the broad scalp topographies of actual EEG. Supplementary Figure 3 depicts the set of 10,000 topographies. Seven topographies were randomly sampled without replacement from the 10,000 surrogates for each of 5,000 permutations. This maintained the dimensionality of the microstate configuration subspace for each permutation. Supplementary Figure 3 also depicts a set of 7 random surrogate topographies. Each set of seven random noise topographies were orthogonalized with SVD and removed from the eyes-closed resting EEG using OSP. The percentage of variance explained by the spatially smoothed noise topographies was obtained for each permutation to construct a surrogate distribution of explained variance for each EEG recording. The specific set of seven maps closest to the mean of each recording's surrogate distribution was selected for subsequent steps. The set of seven representative surrogates was removed from the EEG using OSP to obtain the residual EEG for each recording. The periodic and aperiodic components of the power spectrum were subsequently parameterized. This set of seven topographies was also source localized to provide equivalent methodological controls for the source-level spectral outcomes.

### Supplementary Results

We parameterized periodic and aperiodic features of the PSD and compared the original EEG to the residual EEG following the targeted removal of subspaces of microstate configurations through OSP.

**Removing the Spatial Projections of Microstates Attenuates Parameterized EEG Spectra**

Statistical comparisons of the parameterized power spectrum confirmed that both the periodic and aperiodic features of the macroscopic EEG depend on the spatial architecture of microstates. The results are organized below according to sensor and source-level OSP and periodic and aperiodic spectral features. Supplementary Figure 4 depicts the PSD across both linear and semi-logarithmic scales for the residual EEG after sensor-level removal of microstate configurations compared to the original EEG. The model-estimated effects of removing specific microstate configurations from the original data are summarized in Supplementary Table 1, Supplementary Table 2, and Supplementary Table 3. The threshold for reporting significant tests of parameter estimates was $p < .001$ to account for multiple comparisons across parameterized features of the residual EEG datasets. We first confirmed acceptable *specparam* model fit for the original EEG and after configurations were removed at the sensor level. The mean estimated $R^2$ of parameterized models was 0.978 (95% CI [0.977, 0.980], range = 0.953 – 0.991) for the original data across recording blocks and 0.976 (95% CI [0.974, 0.977], range = 0.930 – 0.988) after sensor-level removal of all microstate configurations.

**Sensor-level OSP on periodic peak alpha and theta power.** We observed a significant main effect of removing specific microstate configurations from the original data on aperiodic-adjusted alpha power, $F(8, 2202) = 400.49$, $p < .001$, and aperiodic-adjusted theta power, $F(8, 1731) = 89.63$, $p < .001$. The residual signal had significantly lower aperiodic-adjusted alpha power ($b = -0.3422$, $SE = 0.0079$, $p < .001$, 95% CI [-0.3577, -0.3267]) and theta power ($b = -0.1164$, $SE = 0.0053$, $p < .001$, 95% CI [-0.1269, -0.1060]) when all configurations were removed, compared to the original EEG (see Supplementary Figure 5). Microstate configurations A, B, C, D, and E, reduced alpha power (*p*s < .001) when separately removed from the original

EEG (see Supplementary Table 1). In contrast, configurations F and G did not significantly change alpha power when removed. Similarly, all microstates except configurations F and G significantly (*ps* < .001) reduced theta power when removed from the original EEG (see Supplementary Table 1). Removing configuration C reduced both alpha and theta power significantly (*p*s < .001) more than all other microstates.

We also observed a significant main effect of sensor-level removal of specific microstate configurations on the alpha center frequency, $F(8, 2236) = 7.53$, $p < .001$, alpha peak bandwidth, $F(8, 2233) = 34.80$, $p < .001$, theta center frequency, $F(8, 1750) = 7.22$, $p < .001$, and theta peak bandwidth, $F(8, 1735) = 23.98$, $p < .001$. The residual signal had significantly lower alpha peak bandwidth ($b = -0.2045$, $SE = 0.0179$, $p < .001$, 95% CI [-0.2395, -0.1694]) and lower theta center frequency ($b = -0.0801$, $SE = 0.0178$, $p < .001$, 95% CI [-0.1151, -0.0452]) when all configurations were removed, compared to the original EEG (see Supplementary Figure 5). In contrast, the removal of all microstate configurations did not significantly (*ps* > .001) affect the alpha center frequency or theta peak bandwidth. The remaining effects of sensor-level removal of specific microstates are provided in Supplementary Table 2.

**Sensor-level OSP on aperiodic 1/*f* offset and exponent.** We observed a significant main effect of sensor-level removal of specific microstate configurations on the aperiodic offset, $F(8, 2256) = 5591.39$, $p < .001$, and aperiodic exponent, $F(8, 2273) = 433.20$, $p < .001$. The residual signal had significantly lower aperiodic offset ($b = -1.1551$, $SE = 0.0060$, $p < .001$, 95% CI [-1.1668, -1.1434]) and lower (flatter) aperiodic exponent ($b = -0.2502$, $SE = 0.0049$, $p < .001$, 95% CI [-0.2599, -0.2405]) when all configurations were removed, compared to the original EEG (see Supplementary Figure 5). The remaining effects of sensor-level removal of specific microstates are provided in Supplementary Table 1. All microstate configurations reduced (*ps* <

.001) the aperiodic offset when removed from the original EEG. All microstates configurations except configuration G reduced (*ps* < .001) the aperiodic exponent. Overall, the targeted removal of microstate configurations at the sensor-level revealed widespread reductions in both the aperiodic offset and exponent for most configurations.

**Source-level OSP on periodic center frequency and bandwidth.** We observed a significant main effect of source-level removal of specific microstate configurations on the alpha center frequency, $F(8, 2236) = 7.53$, $p < .001$, alpha peak bandwidth, $F(8, 2233) = 34.80$, $p < .001$, theta center frequency, $F(8, 1746) = 3.76$, $p < .001$, and theta peak bandwidth, $F(8, 1739) = 22.82$, $p < .001$. The residual signal had significantly lower alpha peak bandwidth ($b = -0.2798$, $SE = 0.0173$, $p < .001$, 95% CI [-0.3137, -0.2459]) and lower theta peak bandwidth ($b = -0.1229$, $SE = 0.0151$, $p < .001$, 95% CI [-0.1526, -0.0932]) when all configurations were removed, compared to the original EEG (see Supplementary Figure 5). In contrast, the removal of all microstate configurations did not significantly (*ps* > .001) affect the alpha center frequency or theta center frequency. The remaining effects of source-level removal of specific microstates are provided in Supplementary Table 3.

**Periodic and aperiodic features of sensor-level removal of surrogate comparison topographies.** We parameterized the PSD of the residual EEG following the sensor-level removal of spatially smoothed surrogate noise topographies to confirm that the attenuation of spectral features was specific to the spatial architecture of microstates. The removal of surrogate topographies at the sensor level resulted in significant but comparatively small reductions to periodic alpha and theta peak power, and aperiodic spectral features (see Supplementary Table 4). However, these effects were substantially smaller in magnitude than those observed following the removal of the microstate configurations. For example, reduction in peak alpha power ($b$ = -

0.0467, $SE = 0.0068$, $p < .001$, 95% CI [-0.0601, -0.0333]) was roughly seven times smaller in magnitude than when the microstate configurations were removed at the sensor-level. Likewise, the reduction in aperiodic exponent ($b = -0.0706$, $SE = 0.0046$, $p < .001$, 95% CI [-0.0795, -0.0616]) was roughly five times smaller than when microstate configurations were removed at the sensor-level.

**Removing the Spatial Projections of Microstates Attenuates Configuration Specific Spectra**

To evaluate the electrode-level effects of sensor-level removal of microstates on the aperiodic-adjusted alpha and theta power, aperiodic offset, and aperiodic exponent for all electrodes in the montage, we compared each electrode from the original continuous EEG time series to the same electrode following the targeted removal of subspaces of microstates through OSP. Maps of the model-estimated effects were thresholded at the Nyholt-corrected significance level to control the family-wise error rate because of tests across the entire 128-channel montage before and after sensor-level removal of microstate configurations. The threshold for significance was $\alpha = 0.001$ for aperiodic-adjusted alpha, $\alpha = 0.00039$ for aperiodic-adjusted theta, $\alpha = 0.00094$ for the aperiodic offset, and $\alpha = 0.00074$ for the aperiodic exponent. Topographical representations of the original, residual, and difference signals are depicted in Supplementary Figures 5 and 6. Together, these results demonstrate that the spatial distribution of spectral changes in the residual signal recapitulates the distinct voltage configurations of the removed microstates.

We observed that removing the spatial projections of all microstates yielded highly significant reductions in alpha power, theta power, aperiodic offset, and aperiodic exponent across all scalp electrodes (see Supplementary Figure 5). The isolated removal of individual microstates instead revealed highly specific spatial effects. The removal of other microstates resulted in localized reductions in alpha power, aperiodic offset, and aperiodic exponent, corresponding precisely to their unique spatial configurations (see Supplementary Figure 6). The removal of microstate C most strongly attenuated the periodic and aperiodic components compared to other microstates. These reductions occurred at anterior and posterior locations aligned with its voltage topography. In contrast, the removal of microstates F and G, which had minimal effect on overall alpha power, demonstrated significant but spatially constrained differences in alpha power, aperiodic offset, and aperiodic exponent.

Supplementary Table 1: Model parameter estimates comparing periodic and aperiodic spectral parameters of the original to the residual EEG after sensor-level removal

| | Aperiodic-adjusted Alpha Power | Aperiodic-adjusted Theta Power | Aperiodic Offset | Aperiodic Exponent |
|---|---|---|---|---|
| Fixed Effects | | | | |
| Intercept (Original EEG) | 1.1269 (0.0426)*** | 0.4436 (0.0245)*** | 0.9517 (0.0347)*** | 1.4208 (0.0207)*** |
| All removed | -0.3422 (0.0079)*** | -0.1164 (0.0053)*** | -1.1551 (0.0060)*** | -0.2502 (0.0049)*** |
| Microstate A removed | -0.0375 (0.0068)*** | -0.0122 (0.0050)* | -0.2044 (0.0050)*** | -0.0256 (0.0044)*** |
| Microstate B removed | -0.0687 (0.0067)*** | -0.0178 (0.0048)*** | -0.2136 (0.0050)*** | -0.0365 (0.0044)*** |
| Microstate C removed | -0.1821 (0.0072)*** | -0.0429 (0.0049)*** | -0.3555 (0.0050)*** | -0.0812 (0.0044)*** |
| Microstate D removed | -0.0704 (0.0069)*** | -0.0261 (0.0049)*** | -0.2557 (0.0049)*** | -0.0756 (0.0044)*** |
| Microstate E removed | -0.1123 (0.0067)*** | -0.0248 (0.0048)*** | -0.2506 (0.0051)*** | -0.0539 (0.0044)*** |
| Microstate F removed | 0.0099 (0.0072) | -0.0069 (0.0054) | -0.1127 (0.0051)*** | -0.0206 (0.0044)*** |
| Microstate G removed | 0.0200 (0.0074)** | 0.0038 (0.0054) | -0.1084 (0.0051)*** | -0.0043 (0.0044) |
| Random Effects | | | | |
| Intercept variance | 0.1398 (0.0226) | 0.0447 (0.0073) | 0.0931 (0.0150) | 0.0326 (0.0053) |
| Residual variance$_{Original}$ | 0.0151 (0.0009) | 0.0062 (0.0004) | 0.0077 (0.0004) | 0.0059 (0.0003) |
| Residual variance$_{All}$ | 0.0237 (0.0014) | 0.0071 (0.0005) | 0.0142 (0.0008) | 0.0093 (0.0005) |
| Residual variance$_{A}$ | 0.0140 (0.0008) | 0.0056 (0.0004) | 0.0075 (0.0004) | 0.0059 (0.0003) |
| Residual variance$_{B}$ | 0.0131 (0.0008) | 0.0050 (0.0003) | 0.0076 (0.0004) | 0.0060 (0.0003) |
| Residual variance$_{C}$ | 0.0170 (0.0010) | 0.0051 (0.0003) | 0.0081 (0.0005) | 0.0061 (0.0004) |
| Residual variance$_{D}$ | 0.0146 (0.0008) | 0.0052 (0.0003) | 0.0072 (0.0004) | 0.0059 (0.0003) |
| Residual variance$_{E}$ | 0.0126 (0.0007) | 0.0050 (0.0003) | 0.0085 (0.0005) | 0.0063 (0.0004) |
| Residual variance$_{F}$ | 0.0173 (0.0010) | 0.0070 (0.0005) | 0.0081 (0.0005) | 0.0063 (0.0004) |
| Residual variance$_{G}$ | 0.0185 (0.0011) | 0.0075 (0.0005) | 0.0081 (0.0005) | 0.0062 (0.0004) |
| *N* | 78 | 76 | 78 | 78 |
| Obs. | 5580 | 4336 | 5580 | 5580 |

Note: Model parameters representing the fixed effects of removing specific microstate configurations from the original data using sensor-level orthogonal subspace projection on dependent measures of aperiodic-adjusted alpha power, theta power, aperiodic offset, and aperiodic exponent. Effects describe the difference between the residual signal and the original EEG (i.e., the intercept) for all microstates and each individual configuration A through G from multilevel mixed effects models. Random intercept variances and residual variances are also provided. Model estimates were based on a sample size of $N = 78$ with 5580 total observations contributing to analyses ($n = 76$ and 4336 observations for theta power). Standard errors are provided in parentheses. The statistical significance of fixed effects is indicated. * $p < .05$, ** $p < .01$, *** $p < .001$

Supplementary Table 2: Model parameter estimates comparing periodic central frequency and bandwidth of the original to the residual EEG after sensor-level removal

| | Alpha CF | Alpha Bandwidth | Theta CF | Theta Bandwidth |
|---|---|---|---|---|
| Fixed Effects | | | | |
| Intercept (Original EEG) | 10.5043 (0.0973)*** | 2.3710 (0.0660)*** | 6.9463 (0.0713)*** | 1.7623 (0.0373)*** |
| All removed | -0.0280 (0.0166) | -0.2045 (0.0179)*** | -0.0801 (0.0178)*** | -0.0269 (0.0156) |
| Microstate A removed | -0.0146 (0.0156) | -0.0492 (0.0175)** | -0.0327 (0.0179) | 0.0272 (0.0152) |
| Microstate B removed | -0.0377 (0.0156)* | -0.0125 (0.0176) | -0.0335 (0.0174) | 0.0113 (0.0151) |
| Microstate C removed | -0.0590 (0.0162)*** | 0.0570 (0.0184)** | -0.0932 (0.0183)*** | 0.1289 (0.0159)*** |
| Microstate D removed | 0.0478 (0.0162)** | 0.0129 (0.0182) | -0.0130 (0.0176) | 0.0019 (0.0159) |
| Microstate E removed | -0.0383 (0.0160)* | 0.0182 (0.0182) | -0.0662 (0.0172)*** | 0.0741 (0.0154)*** |
| Microstate F removed | 0.0101 (0.0162) | -0.0269 (0.0186) | -0.0120 (0.0186) | -0.0426 (0.0162)** |
| Microstate G removed | -0.0138 (0.0165) | -0.0753 (0.0184)*** | -0.0054 (0.0179) | -0.0387 (0.0158)* |
| Random Effects | | | | |
| Intercept variance | 0.7280 (0.1175) | 0.3275 (0.0530) | 0.3723 (0.0617) | 0.0953 (0.0160) |
| Residual variance$_{Original}$ | 0.0800 (0.0046) | 0.0988 (0.0057) | 0.0748 (0.0049) | 0.0564 (0.0037) |
| Residual variance$_{All}$ | 0.0898 (0.0053) | 0.0990 (0.0059) | 0.0740 (0.0050) | 0.0570 (0.0039) |
| Residual variance$_{A}$ | 0.0711 (0.0041) | 0.0913 (0.0052) | 0.0790 (0.0051) | 0.0536 (0.0035) |
| Residual variance$_{B}$ | 0.0716 (0.0041) | 0.0935 (0.0054) | 0.0717 (0.0046) | 0.0541 (0.0035) |
| Residual variance$_{C}$ | 0.0836 (0.0049) | 0.1102 (0.0066) | 0.0865 (0.0057) | 0.0651 (0.0043) |
| Residual variance$_{D}$ | 0.0824 (0.0048) | 0.1070 (0.0062) | 0.0723 (0.0047) | 0.0640 (0.0042) |
| Residual variance$_{E}$ | 0.0784 (0.0046) | 0.1055 (0.0061) | 0.0677 (0.0044) | 0.0581 (0.0038) |
| Residual variance$_{F}$ | 0.0827 (0.0048) | 0.1157 (0.0067) | 0.0855 (0.0058) | 0.0651 (0.0044) |
| Residual variance$_{G}$ | 0.0888 (0.0051) | 0.1119 (0.0065) | 0.0760 (0.0050) | 0.0603 (0.0040) |
| *N* | 78 | 78 | 76 | 76 |
| Obs. | 5580 | 5580 | 4336 | 4336 |

Note: Model parameters representing the fixed effects of removing specific microstate configurations from the original data using sensor-level orthogonal subspace projection on dependent measures of periodic central frequency and bandwidth. Effects describe the difference between the residual signal and the original EEG (i.e., the intercept) for all microstates and each individual configuration A through G from multilevel mixed effects models. Random intercept variances and residual variances are also provided. Model estimates were based on a sample size of $N = 78$ with 5580 total observations contributing to analyses ($n = 76$ and 4336 observations for theta central frequency and bandwidth). Standard errors are provided in parentheses. The statistical significance of fixed effects is indicated. * $p < .05$, ** $p < .01$, *** $p < .001$

Supplementary Table 3: Model parameter estimates comparing periodic central frequency and bandwidth of the original to the residual EEG after source-level removal

| | Alpha CF | Alpha Bandwidth | Theta CF | Theta Bandwidth |
|---|---|---|---|---|
| Fixed Effects | | | | |
| Intercept (Original EEG) | 10.5044 (0.0980)*** | 2.3773 (0.0649)*** | 6.9266 (0.0747)*** | 1.7588 (0.0362)*** |
| All microstates removed | -0.0068 (0.0166) | -0.2798 (0.0173)*** | -0.0287 (0.0183) | -0.1229 (0.0151)*** |
| Microstate A removed | -0.0140 (0.0155) | -0.0936 (0.0173)*** | -0.0254 (0.0178) | -0.0023 (0.0150) |
| Microstate B removed | -0.0352 (0.0157)* | -0.0680 (0.0174)*** | -0.0357 (0.0179)* | -0.0236 (0.0150) |
| Microstate C removed | -0.0652 (0.0167)*** | -0.0142 (0.0182) | -0.0740 (0.0191)*** | 0.0471 (0.0150)** |
| Microstate D removed | 0.0495 (0.0163)** | -0.0215 (0.0181) | 0.0062 (0.0182) | -0.0372 (0.0156)* |
| Microstate E removed | -0.0415 (0.0159)** | -0.0300 (0.0185) | -0.0436 (0.0177)* | 0.0359 (0.0160)* |
| Microstate F removed | 0.0198 (0.0159) | -0.0511 (0.0185)** | -0.0068 (0.0183) | -0.0618 (0.0161)*** |
| Microstate G removed | -0.0040 (0.0160) | -0.0906 (0.0180)*** | 0.0030 (0.0180) | -0.0500 (0.0155)** |
| Random Effects | | | | |
| Intercept variance | 0.7395 (0.1194) | 0.3166 (0.0513) | 0.4156 (0.0686) | 0.0909 (0.0152) |
| Residual variance$_{Original}$ | 0.0796 (0.0046) | 0.0986 (0.0057) | 0.0732 (0.0048) | 0.0534 (0.0035) |
| Residual variance$_{All}$ | 0.0917 (0.0054) | 0.0866 (0.0051) | 0.0808 (0.0055) | 0.0518 (0.0035) |
| Residual variance$_{A}$ | 0.0697 (0.0040) | 0.0871 (0.0050) | 0.0775 (0.0050) | 0.0543 (0.0035) |
| Residual variance$_{B}$ | 0.0740 (0.0043) | 0.0895 (0.0052) | 0.0813 (0.0052) | 0.0549 (0.0035) |
| Residual variance$_{C}$ | 0.0935 (0.0055) | 0.1061 (0.0063) | 0.1029 (0.0067) | 0.0545 (0.0036) |
| Residual variance$_{D}$ | 0.0854 (0.0050) | 0.1037 (0.0060) | 0.0811 (0.0054) | 0.0605 (0.0040) |
| Residual variance$_{E}$ | 0.0780 (0.0046) | 0.1146 (0.0067) | 0.0766 (0.0050) | 0.0690 (0.0045) |
| Residual variance$_{F}$ | 0.0764 (0.0044) | 0.1125 (0.0065) | 0.0837 (0.0055) | 0.0675 (0.0045) |
| Residual variance$_{G}$ | 0.0796 (0.0046) | 0.1026 (0.0059) | 0.0784 (0.0052) | 0.0587 (0.0039) |
| *N* | 78 | 78 | 77 | 77 |
| Obs. | 5580 | 5580 | 4317 | 4317 |

Note: Model parameters representing the fixed effects of removing specific microstate configurations from the original data using source-level orthogonal subspace projection on dependent measures of periodic central frequency and bandwidth. Effects describe the difference between the residual signal and the original EEG (i.e., the intercept) for all microstates and each individual configuration A through G from multilevel mixed effects models. Random intercept variances and residual variances are also provided. Model estimates were based on a sample size of $N = 78$ with 5580 total observations contributing to analyses ($n = 77$ and 4317 observations for theta central frequency and bandwidth). Standard errors are provided in parentheses. The statistical significance of fixed effects is indicated. * $p < .05$, ** $p < .01$, *** $p < .001$

Supplementary Table 4: Model parameter estimates comparing spectral parameters of the original EEG to the smoothed noise surrogates after sensor-level removal

| | Aperiodic-adjusted Alpha Power | Aperiodic-adjusted Theta Power | Aperiodic Offset | Aperiodic Exponent |
|---|---|---|---|---|
| Intercept (Original EEG) | 1.1269 (0.0449)*** | 0.4482 (0.0262)*** | 0.9517 (0.0350)*** | 1.4209 (0.0213)*** |
| Smoothed noise removed | -0.0467 (0.0068)*** | -0.0247 (0.0050)*** | -0.4209 (0.0052)*** | -0.0706 (0.0046)*** |
| Random Effects | | | | |
| Intercept variance | 0.1554 (0.0252) | 0.0491 (0.0083) | 0.0947 (0.0153) | 0.0346 (0.0056) |
| Residual variance$_{Original}$ | 0.0141 (0.0008) | 0.0062 (0.0004) | 0.0078 (0.0005) | 0.0061 (0.0004) |
| Residual variance$_{Noise}$ | 0.0148 (0.0009) | 0.0055 (0.0004) | 0.0087 (0.0005) | 0.0068 (0.0004) |
| *N* | 78 | 73 | 78 | 78 |
| Obs. | 1240 | 936 | 1240 | 1240 |

Note: Model parameters representing the fixed effects of removing the smoothed noise surrogates from the original data using sensor-level orthogonal subspace projection on dependent measures of aperiodic-adjusted alpha power, theta power, aperiodic offset, and aperiodic exponent. Effects describe the difference between the residual signal and the original EEG (i.e., the intercept) from multilevel mixed effects models. Random intercept variances and residual variances are also provided. Model estimates were based on a sample size of $N = 78$ with 1240 total observations contributing to analyses ($n = 73$ and 936 observations for theta power). Standard errors are provided in parentheses. The statistical significance of fixed effects is indicated. * $p < .05$, ** $p < .01$, *** $p < .001$

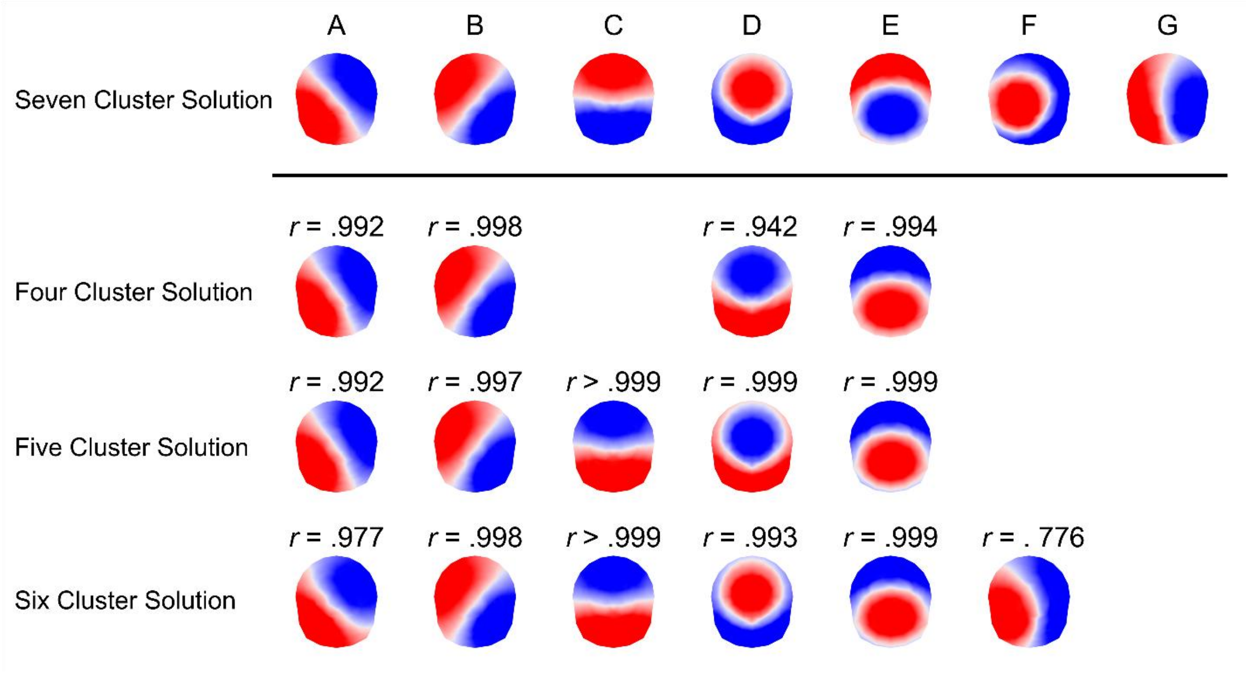


Supplementary Figure 1: Topographic correlations are depicted between the configurations of the seven-cluster microstate solution and lower-order cluster solutions. The top row displays the seven global microstate configurations (A–G) utilized in the primary analyses. The subsequent rows display the topographic configurations identified when *k*-means clustering was constrained to four-, five-, and six-cluster solutions. Each lower-order topography is aligned beneath its counterpart in the seven-cluster solution. Spatial correlation coefficients (*r*) are provided above each configuration with the corresponding seven-cluster configuration above. Correlation values were based on the relative topographical configuration but not the polarity of the topographies. The spatial correlations demonstrate that the topographies of lower-order solutions are encompassed within the spatial subspaces defined by the seven-cluster repertoire.

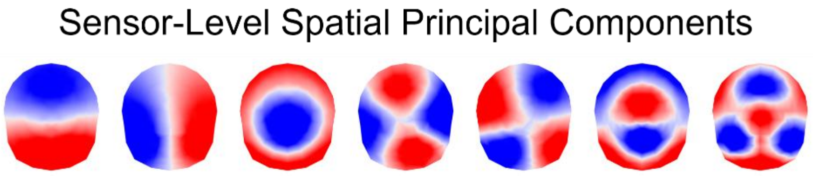


Supplementary Figure 2: Grand-average spatial components derived from principal components analysis (PCA) of the 128-channel EEG. The topographies represent the eigenvectors of the first seven principal components calculated from the mean spatial covariance matrix of the 128-channel EEG recordings across all recordings. The seven spatial components explained 95.03% (95% CI [94.59, 95.46]) of variance. The first principal component (leftmost map) has a near-perfect spatial correlation ($r$ = .998) with the topography of microstate configuration C.

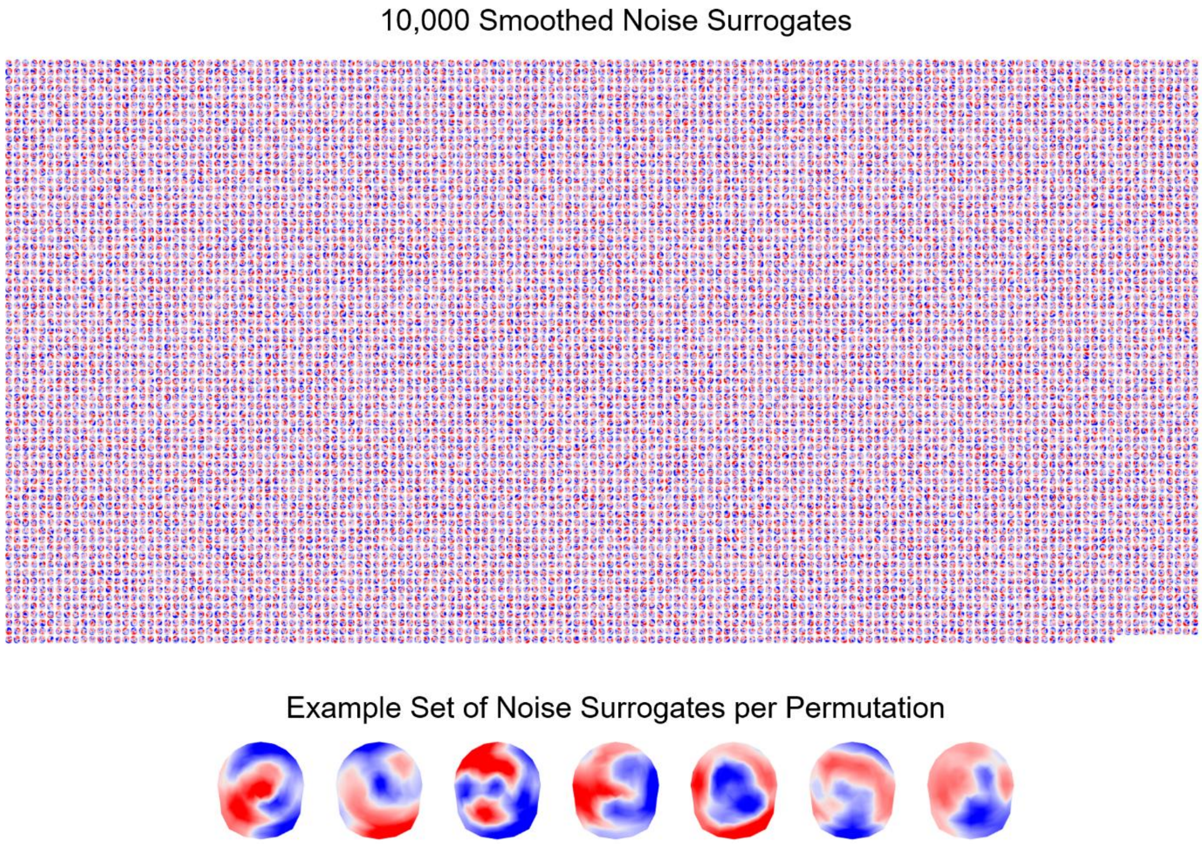


Supplementary Figure 3: A pool of 10,000 iteratively smoothed noise surrogates was generated as surrogate control topographies for subspace projections. The pool of 10,000 smoothed noise surrogates is shown at the top. Surrogates were constructed to resemble the spatial autocorrelation and smoothness of real EEG. Random sets of seven maps (bottom) were iteratively removed via OSP to calculate explained variance and construct a null distribution of 5,000 permutations characterizing the effects of removing smoothed noise on the EEG. The specific set of seven maps closest to the mean of each recording's distribution was selected for subsequent steps. The set of seven representative surrogates was removed from the EEG using OSP to obtain the residual EEG for each recording. The periodic and aperiodic components of the power spectrum were subsequently parameterized. This set of seven topographies was also source localized to provide equivalent methodological controls for the source-level spectral outcomes.

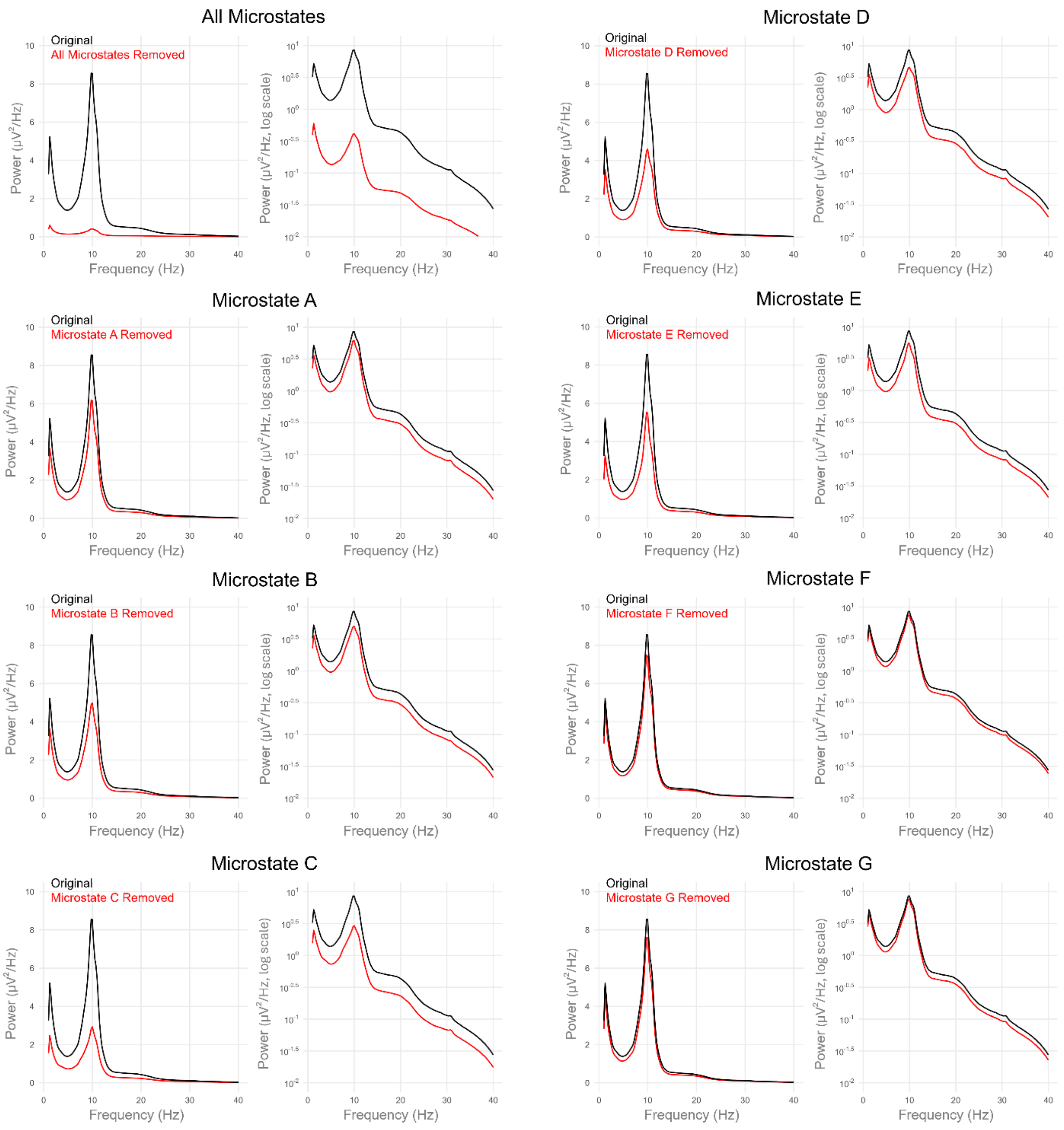


Supplementary Figure 4: Power spectral density (PSD) of the original continuous EEG is shown compared to the residual signal following the computational removal of microstate topographies at the sensor level using orthogonal subspace projection (OSP). Each panel displays the PSD across the frequency range on both a linear scale (left) to emphasize the periodic oscillatory peaks and a semi-logarithmic scale (right) to emphasize the broadband aperiodic 1/*f* background. The intact EEG signal is depicted in black, while the residual signal is depicted in red following OSP of all seven microstates combined (top left) or the selective removal of individual microstate configurations A through G.

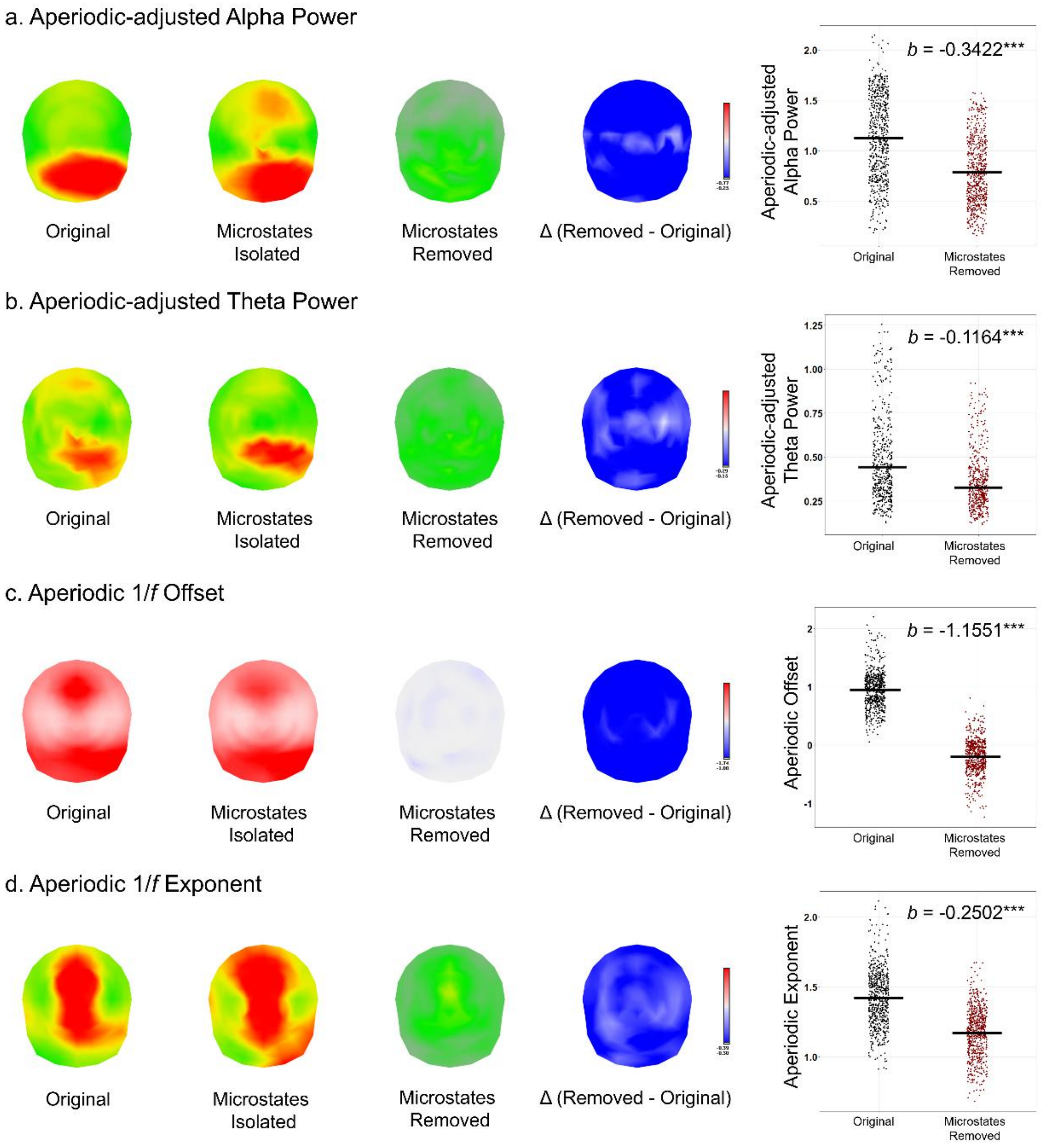


Supplementary Figure 5: Topographic maps illustrate the grand average topography of the intact EEG signal (original), the grand average signal reconstructed from the sensor-level spatial projections of microstates (microstates isolated), the grand average residual signal after sensor-level computational removal all microstates (microstates removed), and the model-estimated difference, Δ (removed – original), for a). the aperiodic-adjusted alpha power, b). the aperiodic-adjusted theta power, c). aperiodic 1/*f* offset, and d). aperiodic 1/*f* exponent. For the difference-maps, decreases are shown in blue at electrode locations and electrode locations with non-significant differences are omitted from the map. Jitter plots (right) display the corresponding individual data points from recording epochs for the original (black) and residual (red) signals averaged across the electrode montage. The horizontal black lines denote the model-estimated mean value. Unstandardized beta coefficients (*b*) denote the model-estimated reduction in each spectral parameter following the removal of all microstates. *** $p < .001$

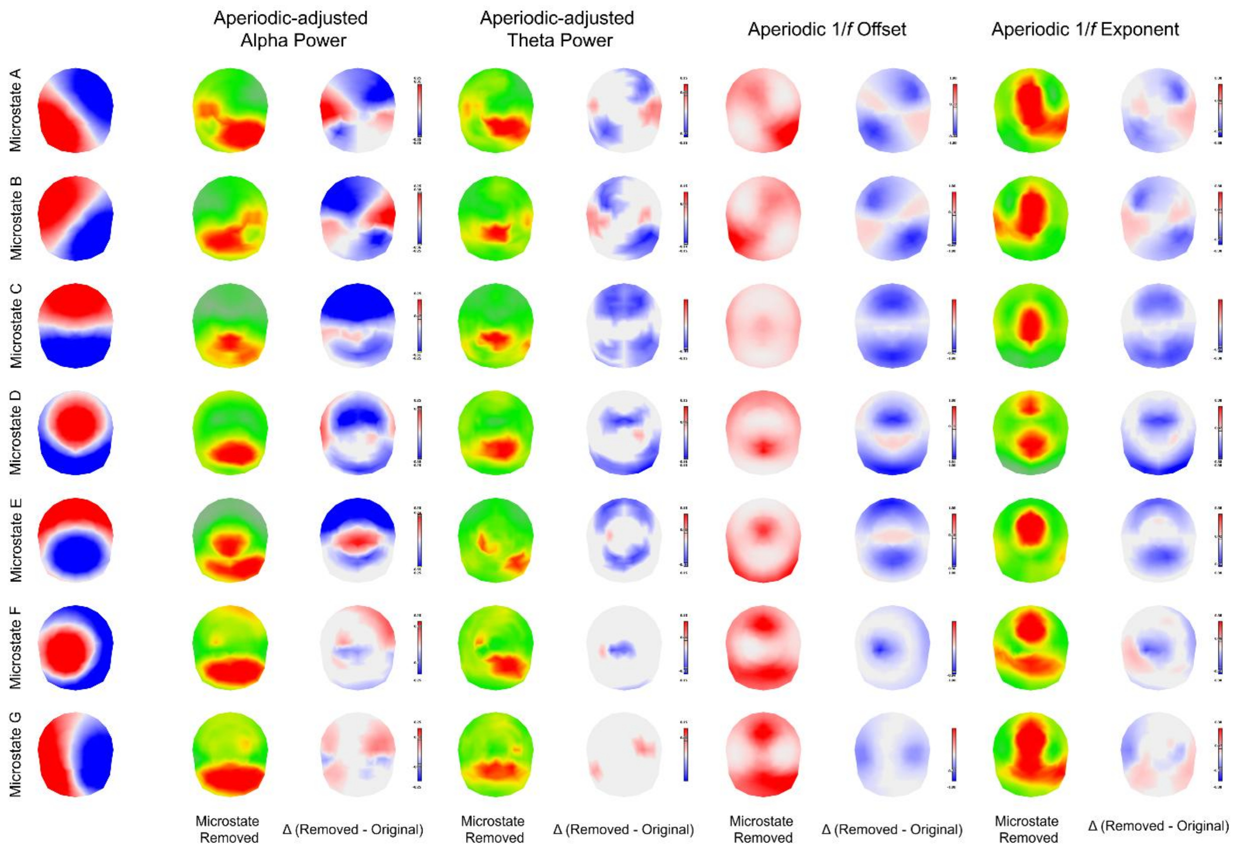


Supplementary Figure 6: Topographic maps illustrating the spatially specific effects of computationally removal individual microstate configurations (A through G) at the sensor level. The leftmost column displays the voltage topography of each targeted microstate. The subsequent columns present the topography of grand average residual signal (microstate removed) and the model estimated difference, Δ (removed – original), for the aperiodic-adjusted alpha power, the aperiodic-adjusted theta power, aperiodic 1/*f* offset, and aperiodic 1/*f* exponent. Across all comparison maps, significant increases are shown in red (indicating greater power at that electrode location when that microstate is removed), while decreases are shown in blue. Electrode locations with non-significant differences are omitted.
